# Long-term co-circulation of multiple arboviruses in southeast Australia revealed by xeno-monitoring and metatranscriptomics

**DOI:** 10.1101/2024.03.29.587110

**Authors:** Carla Julia S. P. Vieira, Michael B. Onn, Martin A. Shivas, Damien Shearman, Jonathan M. Darbro, Melissa Graham, Lucas Freitas, Andrew F. van den Hurk, Francesca D. Frentiu, Gabriel L. Wallau, Gregor J. Devine

## Abstract

Arbovirus surveillance of wild-caught mosquitoes is an affordable and sensitive means of monitoring virus transmission dynamics at various spatial-temporal scales, and emergence and re-emergence during epidemic and interepidemic periods. A variety of molecular diagnostics for arbovirus screening of mosquitoes (known as xeno-monitoring) are available, but most provide limited information about virus diversity. PCR-based screening coupled with metatranscriptomics is an increasingly affordable and sensitive pipeline for integrating complete viral genome sequencing into surveillance programs. This enables large-scale, high-throughput arbovirus screening from diverse samples. We collected mosquitoes in CO2-baited light traps from five urban parks in Brisbane from March 2021 to May 2022. Mosquito pools of ≤200 specimens were screened for alphaviruses and flaviviruses using virus genus-specific primers and reverse transcription quantitative PCR (qRT-PCR). A subset of virus-positive samples was then processed using a mosquito-specific ribosomal RNA depletion method and then sequenced on the Illumina NextSeq. Overall, 54,670 mosquitoes, representing 26 species were screened in 382 pools. Thirty detections of arboviruses were made in 28 pools. Twenty of these positive pools were further characterised using meta-transcriptomics generating 18 full-length genomes. These full-length sequences belonged to four medically relevant arboviruses: Barmah Forest, Ross River, Sindbis-like and Stratford viruses. Phylogenetic and evolutionary analyses revealed the evolutionary progression of arbovirus lineages over the last 100 years, highlighting long-distance dispersal across the Australian continent and continuous circulation characterised by constant turnover of virus lineages.

## 1. Introduction

In recent decades, arthropod-borne viruses (arboviruses) have emerged or re-emerged as important human and animal pathogens with tremendous implications for public health globally (Huang et al., 2019). Dengue viruses (DENV) alone infect around 390 million people annually (WHO, 2023) and are associated with a substantial economic burden (US$40 billion) (Selck et al., 2014; Stanaway et al., 2016). Although many arboviruses were geographically restricted, the continued movement of people and vectors around the globe have expanded their ranges (Huang et al., 2019). For example, DENV and Chikungunya (CHIKV), both emerging in Africa, have now spread to every continent except Antarctica (Gubler, 1998; Nunes et al., 2015) while Japanese encephalitis virus (JEV) has been expanding its range in Asia, and has recently become endemic across Australia (Yakob et al., 2023). Climate heating, extreme weather events and anthropogenic factors such as land-use, urbanisation and globalisation facilitate that invasion process by mediating changes in vector and reservoir distributions, and increasing immigration of viraemic hosts (Patz and Norris, 2004; Vieira et al., 2020).

In Australia, endemic arboviruses cause a significant health and economic burden. The include the arthritogenic alphaviruses Ross River virus (RRV) and Barmah Forest virus (BFV), which respectively cause approximately 5,000 and 1,000 cases annually, and inducing significant health impacts (Department of Health and Aged Care, 2023). These alphaviruses are also endemic in Papua New Guinea (PNG) (Kizu et al., 2023) while RRV has spread across several Pacific Island Countries and Territories (PICTs) where it caused an epidemic of an estimated 500,000 cases in 1979–1980 (Marshall and Miles, 1984). RRV has the potential for global spread (Lau et al., 2017; Shanks, 2019). Additionally, Australia experiences sporadic outbreaks of encephalitogenic flaviviruses, including the Kunjin strain of West Nile virus (WNVKUN), Murray Valley encephalitis virus (MVEV), and JEV, with mortality rates of 15% for the 2022 JEV outbreak and 23% for the 2023 MVE outbreak (Department of Health and Aged Care, 2023). Over 75 other alpha- and flaviviruses are endemic in Australia (CDC, 2023), some with the potential to cause symptoms in humans. Their public health significance remains poorly understood as do their key vectors, reservoirs and spill-over risks (Gyawali et al., 2019).

Most arboviruses that infect humans belong to the *Alphavirus* and *Flavivirus* genera, in the Togaviridae and Flaviviridae families, respectively. The majority of human infections by these viruses are asymptomatic or subclinical meaning that the emergence or re-emergence of arboviruses may go undetected and unreported (Grubaugh et al., 2019a). A salient case is cryptic circulation of the ZIKV in Brazil for more than 18 months before its first detection and facilitating its spread to over 40 countries (Faria et al., 2017; Grubaugh et al., 2019b). Another example in the Australian context is the recent JEV range expansion from Indonesia to Australia. The genotype of JEV currently circulating in Australia was detected in the Tiwi Islands, Northern Territory (NT), in February 2021, and one year later in other states of Australia (Sikazwe et al., 2022). However, it is estimated that the Australian clade emerged six years ago, suggesting that this clade of GIV viruses had been cryptically circulating in Australia for several years (95% HPD, 2–14) (Xu et al., 2023). There were 45 human cases of this vaccine-preventable disease recorded during 2022, including seven deaths. It also caused significant stock losses in piggeries (Zhang et al., 2023). It is possible that detrimental impacts of ZIKV in Brazil and JEV in Australia could have been reduced had there been an effective continuous surveillance for early detection and mitigation of these arboviruses, particularly with regard to vector control measures for *Aedes aegypti* (Ritchie et al., 2021) and the provision and targeting of JEV vaccines in humans (Furuya-kanamori et al., 2022).

Mosquito surveillance programs can play a pivotal role in public health strategy and disease preparedness. Xeno-monitoring – the detection of pathogens in arthropod vectors – is a widely used technique for (1) monitoring the distribution of existing and emerging pathogens (Cameron and Ramesh, 2021) and 2) characterising the risks and pathways associated with their transmission to humans. Xeno-monitoring of arboviruses from field collected mosquitoes can be conducted using various methods, including (1) virus culture, (2) PCR-based detection, e.g. quantitative real-time reverse transcription PCR (qRT-PCR), and, more recently, (3) sequencing, e.g. next-generation sequencing (NSG). These techniques have been extensively employed to map an enormous diversity of arboviruses across different mosquito populations and environments (Moonen et al., 2023). The NGS approaches, e.g. metatranscriptomics (total RNA sequencing), facilitates the non-targeted, high-throughput detection and characterisation of whole arbovirus genomes (Batovska et al., 2019; Pronyk et al., 2023). This powerful tool can assist in the characterisation of novel or emerging pathogens or yield genome-wide information that increases our understanding of the transmission dynamics of different virus genomes in different environments and hosts (Batovska et al., 2018; Young et al., 2020).

In Australia, besides monitoring human cases, most states and territories conduct active arbovirus surveillance. This includes seasonally screening mosquitoes for specific arboviruses that are nationally notifiable, using virus isolation and/or qRT-PCR techniques (Knope et al., 2019). Sequencing is not a routine part of the surveillance, so, despite the wide distribution and clinical significance of several arboviruses, whole genome sequences have only been recorded in a few studies (Michie et al., 2023, 2021, 2020). Although NGS is widely available in developed countries (Pronyk et al., 2023), genomic surveillance networks for zoonotic pathogens other than SARS-CoV-2 are still in their early stages. They can be challenging to implement due to their technical complexity, costs and scalability. To address this issue, we examined the use of xeno-diagnostic tools (qRT-PCR) and metatranscriptomics to characterise arboviruses present in wild mosquito populations in Brisbane, the state capital of Queensland, Australia. Our results reveal co-circulation of multiple arboviruses maintained by a wide range of vector species. Some of these are of considerable importance in a public health context. We also discuss how the monitoring approaches that we used might augment existing disease surveillance programs.

## 2. Methods

### 2.1. Study area

This study was carried out in Brisbane, Queensland (27°28’12’’ S and 153°01’15’’ E) between March 2021 and May 2022. Brisbane is the third most populous city and the largest state capital in Australia by geographic area. It has a sub-tropical climate, a rainy season from November to March (annual precipitation levels of 1011.5 mm) and monthly average temperatures of 10–22°C and 20–29°C in winter and summer, respectively (BoM, 2023). The greater Brisbane area contains many habitat types including freshwater, estuarine wetlands, saltmarshes, mangroves, bushlands and subtropical rainforests (Queensland Government, 2023). Brisbane is Australia’s most biodiverse capital city with a variety of native and introduced wildlife, and arbovirus vector and reservoir species. In combination with the high human notification rates for BFV and RRV (Knope et al., 2019), this makes the city an ideal location to investigate endemic and emerging arboviruses that circulate in sylvatic/urban interfaces.

### 2.2. Study design

In Queensland, mosquito-based surveillance systems were initially developed to replace sentinel animals in the remote Torres Strait (Ritchie et al., 2007) but high mosquito trap rates (>100,000 mosquitoes per week) and the logistical challenges of collecting mosquitoes at intervals short enough to preserve viral RNA made processing time-consuming, labour-intensive, and inefficient (van den Hurk et al., 2012). To address these challenges, sugar-based surveillance systems have been developed to detect pathogens in the saliva of infected mosquitoes (Hall-Mendelin et al., 2010; van den Hurk et al., 2014). These systems use sugar-baited filter papers that preserve nucleic acids (e.g., Flinders Technology Associates (FTA) cards®). The presence of these in the trap encourages mosquito feeding and expectoration, and the traps are also baited with CO2 to increase catch numbers. This system is currently utilised as part of a limited mosquito and arbovirus surveillance program, funded by the state’s health department (Queensland Health (QH)) with major contributions from some local councils.

Our study was conducted in partnership with Brisbane City Council (BCC) and QH. BCC routinely deploys mosquito traps (light traps) containing honey-soaked FTA cards on a weekly basis, from October to May each year. The FTA cards are screened for two notifiable arboviruses by QH (RRV, BFV and, following the 2022 JEV outbreak). In this study we were notified of positive FTA cards. We then retrieved the corresponding mosquito collections from BCC and, with QH, conducted one additional mosquito trappings per positive FTA card at the same sites, with the aim of increasing the number of virus-infected mosquito pools for testing.

### 2.3. Mosquito collection and identification

Five BCC trap sites representing a diversity of mosquito habitats were chosen. All were urban parks near residential areas. Three sites were within 500 m of a saltmarsh, and the other two sites were close to freshwater habitats (**Table 1**, **Figure 1**). CDC-style light traps (Pacific Biologics, Scarborough, Australia) were baited with 2kg of dry ice as a CO2 source. Traps deployed by BCC were also baited with 1-octen-3-ol (Merck Life Science, Bayswater, Australia) (van Essen et al., 1994). Traps were set once a week prior to dusk and then collected the following morning. The mosquitoes from each trap were transported to the Mosquito Control Laboratory (MCL) where they were cold-anesthetised, identified using dichotomous keys (Marks, 1982; Russell, 1993; Russell and Debenham, 1996) and pooled (up to 200 individuals) according to species, site and collection date. Mosquito pools were stored at −80°C until RNA extraction.

**Figure 1.**
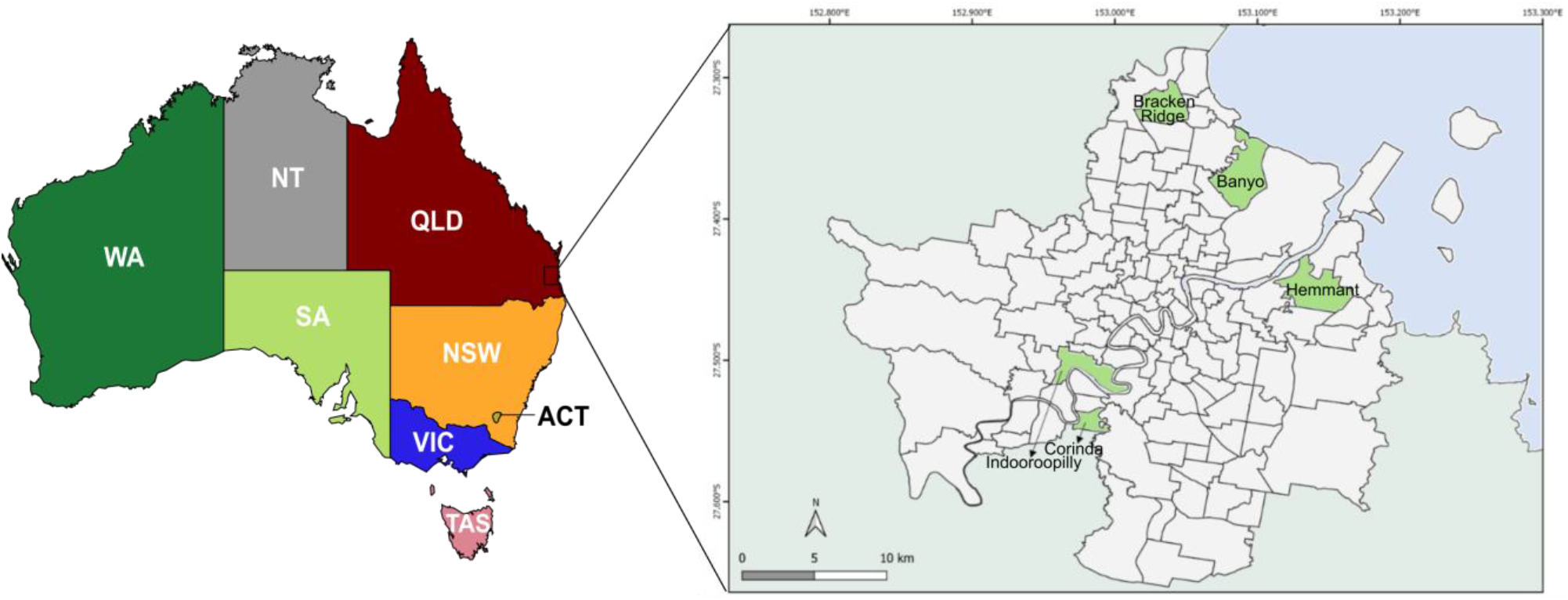
Map of Australia and study area in southeast Queensland. On left, the eight Australian states and territories are shown, colour-coded in accordance with the phylogenetic analysis carried out in this study (see Figure 2). Inset, the five suburbs containing the collection sites are marked in green. Australian states: ACT – Australian Capital Territory; NSW – New South Wales; NT – Northern Territory; QLD – Queensland; SA – South Australia; TAS – Tasmania; VIC – Victoria; WA – Western Australia.

**Table 1.**
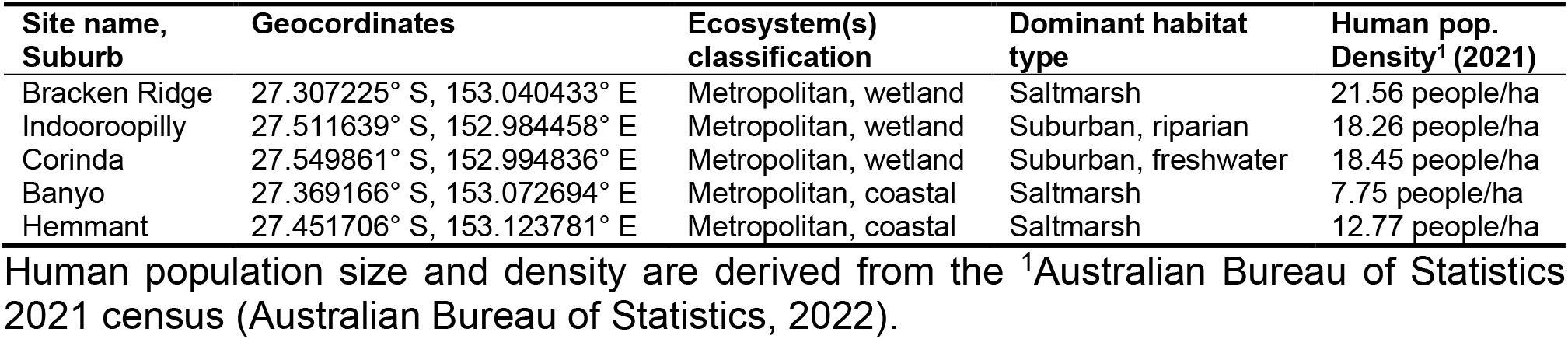
Mosquito trap site location, type, and social characteristics of Brisbane local government area, Queensland, Australia

| Site name, Suburb | Geocoordinates | Ecosystem(s) classification | Dominant habitat type | Human pop. Density<sup>1</sup> (2021) |
| --- | --- | --- | --- | --- |
| Bracken Ridge | 27.307225° S, 153.040433° E | Metropolitan, wetland | Saltmarsh | 21.56 people/ha |
| Indooroopilly | 27.511639° S, 152.984458° E | Metropolitan, wetland | Suburban, riparian | 18.26 people/ha |
| Corinda | 27.549861° S, 152.994836° E | Metropolitan, wetland | Suburban, freshwater | 18.45 people/ha |
| Banyo | 27.369166° S, 153.072694° E | Metropolitan, coastal | Saltmarsh | 7.75 people/ha |
| Hemmant | 27.451706° S, 153.123781° E | Metropolitan, coastal | Saltmarsh | 12.77 people/ha |

Human population size and density are derived from the <sup>1</sup>Australian Bureau of Statistics 2021 census (Australian Bureau of Statistics, 2022).

### 2.4. Sample preparation and nucleic acid extraction

Mosquito pools were homogenised in 2.0 ml Eppendorf Safelock microcentrifuge tubes using DNA/RNA Shield storage buffer (Zymo Research, Irvine, USA) and 2.3 mm zirconium silica beads (Daintree Scientific, St Helens, Australia) scaled according to the mosquito numbers present in each sample (volumes presented in **Table S1**). Mosquito samples were mechanically homogenised for two 3-minute cycles at 1,500 rpm using a Mini Beadbeater-96 (BioSpec Products, Bartlesville, USA) and centrifuged for two 5-minute cycles at 14,000 rpm, at 4°C. Nucleic acid was extracted from the supernatant using a QIAamp Viral RNA Mini Kit (Qiagen, Hilden, Germany) following the manufacturer’s instructions with modifications. The carrier RNA was not added, and a double elution of 40 µL each was performed using UltraPure water (Invitrogen, Carlsbad, USA). An extraction negative control using DNA/RNA Shield was included in each batch of extractions. Extracted nucleic RNA was stored at -80°C until further processing.

### 2.5. Viral detection

Pools were tested for the presence of flaviviruses and alphaviruses using qRT-PCR with genus-specific primers targeting the NS5 and nsP4 regions, following Vina-Rodriguez et al., 2017 and Hermanns et al., 2017 protocols, respectively, with modifications outlined below. Briefly, a SYBR green I based qRT-PCR kit (Luna Universal One-Step NEB, Ipswich, USA) was used in PCR mixtures (20 µl) containing 2 µL cDNA, 10 µL 2X Luna Universal One Step master mix, 0.8 µl WarmStart Luna RT, 5.6 µL UltraPure water, and 0.4 µM (final concentration) of forward and reverse primers designed to either amplify flaviviruses – Pflav-fAAR (TACAACATGATGGGAAAGAGAGAGAA RAA) and PflavrKR (GTGTCCCAKCCRGCTGTGTCATC) 243 bp region – or to amplify alphaviruses – Pan-Alpha-F1 (TCAGCAGAAGAYTTYGAYGC) and Pan-Alpha-R2 (ACATTCCAGAYTTCATCAT) 253 bp region. Thermocycling consisted of 1 cycle of reverse transcription at 55°C for 10 minutes, followed by RT inactivation/Polymerase Activation at 95°C for 1 minute, then 40 cycles of amplification at 95°C for 10 seconds, 50°C for 20 seconds, and 60°C for 30 seconds (data collection). PCR amplification was carried out using the MIC platform (Biomolecular Systems, Sydney, Australia).

For both primer sets, a negative extraction control (see section 2.4), a no template control, and serial 10-fold dilutions of known concentrations of positive controls (CHIKV for alphavirus tests and DENV for flavivirus tests – kindly provided by Dr. Narayan Gyawali, QIMR Berghofer (QIMRB)) were included for melting curve analysis. Amplicons with well-defined melting curve peaks matching the expected range for different alpha- or flaviviruses (*n* = 30 amplicons from 28 samples) were sent to the QIMRB’s Analytical Facility and confirmed by Sanger sequencing (ABI-PRISM 3130 Genetic Analyser, Applied Biosystems, Foster City, USA). Sequences were compared with published sequences using Basic Local Alignment Search Tool and the GenBank database to confirm the identity of the virus. A subset of samples returned detections of medically important arboviruses (*n* = 20 samples) and was then submitted for library preparation and metatranscriptomic sequencing to obtain full genomes. The remaining samples (*n* = 8) returned hits against either insect-specific viruses or unclassified viruses and were not further investigated in this study.

### 2.6. Library preparation and metatranscriptomic sequencing

Extracted samples were DNase treated (NEB, Ipswich, USA) to eliminate any residual DNA molecules from the host in the RNA extracts. After DNase treatment, samples were assessed for RNA quality using the Agilent 2200 TapeStation system (Agilent Technologies, Santa Clara, USA) and subsequently quantified with the Qubit 3.0 fluorometer (Qubit RNA HS and dsDNA HS assay kits; Thermo Fisher Scientific, Waltham, USA), to standardise RNA for Illumina library preparation. All samples had an RNA integrity number (RIN) of >3.0. In the library preparation, a library negative control was used, comprising 20 µL of UltraPure water (Invitrogen, Waltham, USA). For a positive control library, 20 µL of RNA from a pool of 1,000 mosquitoes containing a single RRV infected mosquito was included, as recommended (Batovska et al., 2019). A total of 20 µL of undiluted DNase treated RNA extracts with the amount of RNA ranging from 50 to 500 ng were used to construct sequencing libraries with the Ovation universal transcriptome sequencing (RNA-Seq) system (NuGEN, San Carlos, CA) with customised mosquito rRNA depletion probes (Batovska et al., 2019). Paired-end (150-bp) sequencing of each library (approx. 20 million reads per library) was then performed on the NextSeq 550 platform (Illumina). All library preparation and sequencing procedures were carried out by the Analytical Facility at QIMRB.

### 2.7. Genome assembly and annotation

Reads were assembled using the ViralFlow pipeline (Dezordi et al., 2022), which is a workflow that performs reference-based genome assembly along with several complementary analyses. Briefly, FastQC was used to assess the quality of Illumina raw reads, and low-quality reads were trimmed and mapped against reference genomes, generating consensus sequences at a minimum depth of coverage of 10X. Consensus sequences of viral genomes were obtained through the Integrated Genome Viewer software (Robinson et al., 2011). The consensus sequences generated in this study were deposited in GenBank under accession numbers PP496979 to PP496996 and the raw NGS reads are available the NCBI Sequence Read Archive (SRA) database under BioProject PRJNA1077787.

### 2.8. Bioinformatic analyses

Coding regions of the complete genomes generated in this study were aligned with all published near-complete genomes (>10 kb). Sequences obtained from public databases that presented duplicate strain names for the same strain or that corresponded to cell passages of the same original isolate were removed. All genomes and data (date of isolation, host and location) were collected from ViPR (Pickett et al., 2012): http://www.viprbrc.org/, accessed on 10 September 2023 (accession numbers and genomic information given in **Table S2**). Sequences that had no data described were manually assigned through literature searches. Multiple sequence alignments were performed using MAFFT v7 (Katoh et al., 2019): http://mafft.cbrc.jp/alignment/software/, and manually edited in AliView v1.26 (Larsson, 2014).

Potential recombination events were investigated using the full set of tests implemented in Recombination Detection Program – RDP v4.101 (Martin et al., 2015) and genomes showing recombination events supported by at least three of the seven methods were removed from further analyses (Langat et al., 2020).

### 2.9. Phylogenetic analyses

The optimal evolutionary model was selected using the Akaike information criterion in Smart Model Selection (SMS) (Lefort et al., 2017) implemented in PhyML v3.0 (Guindon et al., 2010) http://www.atgc-montpellier.fr/phyml/. Tamura-Nei (TN93) with gamma distributions (+G) (TN93 + G) gave the best fit for Stratford virus (STRV), general time-reversible (GTR) model with gamma distributions (GTR + G) for BFV and Sindbis-like virus (SINV), and with invariant sites (+I) (GTR + G + I) for RRV. Maximum-likelihood (ML) phylogenies were generated using PhyML v3.0, employing a Subtree Pruning and Regrafting (SPR) topology searching algorithm. We assessed statistical support for phylogenetic branching points using the approximate likelihood ratio test on the Shimodaira-Hasegawa-like procedure (SH-aLRT) with 1,000 replicates.

### 2.10. Molecular clock

To evaluate if the data were appropriate for the estimation of temporal parameters, a regression analysis of the root-to-tip divergence against tip sampling time of the ML phylogenetic trees was performed in TempEst v1.5.3 (Rambaut et al., 2016) (**Table S3**). Outlier sequences that deviated by > 1.5 interquartile ranges from the root-to-tip regression line were excluded. We found strong temporal signals (correlation coefficients ranging from 0.8623 to 0.9758 and R^2^ ranging from 0.6404 to 0.9521), suggesting that the different datasets were appropriate for the estimation of temporal parameters. Bayesian molecular clock phylogenetic analysis was performed for each virus with BEAST 1.10.4 (Suchard et al., 2018) in three independent runs of 100 million Markov chain Monte Carlo (MCMC), sampling every 10,000 generations. After removing 10% burn-in, the final Bayesian consensus tree datasets were generated. The molecular clock and demographic model were chosen by identifying the optimal likelihood combination through Path Sampling/Stepping-stone sampling. This comparison involved evaluating strict and uncorrelated lognormal clock models, each associated with two distinct demographic models: Constant and Bayesian Skyline priors (**Table S4**). Convergence between runs was evaluated with Tracer 1.7.1 (Rambaut et al., 2018) and the final combined dataset showed an effective sample size (ESS) > 200 for all parameters sampled. The tree files of each virus were combined using LogCombiner v1.10.4 and maximum clade credibility (MCC) trees were extracted and summarised using TreeAnnotator v1.10.4. Tree visualisation and figure generation were performed with FigTree v1.4.4 (Rambaut, 2014). SPREAD4 software (Nahata et al., 2022) (https://spreadviz.org/) was used to create geographic maps that allowed the visualisation of discrete and continuous spatio-temporal reconstructions and geographic migration history.

## 3. Results

### 3.1. Mosquito diversity

We were notified of 11 traps collected between March 2021 and May 2022 in our 5 sampling areas that had FTA cards positive for RRV. We recovered the contents of those traps and deployed a total of 11 additional samplings at the same sites, totalling 22 traps. A total of 54,670 adult mosquitoes from 26 species were collected. Species abundance and diversity differed between sites, with greatest overall diversity of species in Banyo (23 species), site with a mix of freshwater, saltmarsh and estuary, compared with the least diverse site, located in Hemmant (7 species), an industrial suburb. Mosquito abundance was also greatest in Banyo (47.4% of all mosquitoes). The dominant species across all sites were *Culex annulirostris* (53.5%), *Cx. orbostiensis* (14.9%), and *Ae. procax* (8.1%) (**Table S5**).

### 3.2. Viral screening and metatranscriptomic sequencing

Mosquitoes were separated into 382 species-specific pools, each pool containing up to 200 mosquitoes. Of these, a total of 30 virus detections was made from 28 pools using qRT-PCR. All detections were confirmed by Sanger sequencing. We detected four medically relevant arboviruses: the alphaviruses BFV, RRV, and Sindbis-like virus (SINV-like) and the flavivirus Stratford virus (STRV). Detections were made from seven mosquito species, mostly *Cx. annulirostris* and *Ae. procax* (**Table 2**).

**Table 2:** Details of viruses detected in mosquitoes collected from Brisbane sites, from March 2021 to May 2022.

| Pool No. | Species | Collection site | Date | QTD | qRT-PCR result / confirmed by Sanger sequencing | RNAseq result | No. total reads | No. mapped reads | % mapped reads | Depth of coverage | Sequence length (nt) | Reference genome | Genome reference length (nt) |
| --- | --- | --- | --- | --- | --- | --- | --- | --- | --- | --- | --- | --- | --- |
| BCC 16 | *Ae. vigilax* | Bracken R | 15/3/2021 | 161 | RRV | RRV | 40,926,873 | 1,003,045 | 2.45 | 6,311 | 11,913 | MH987781.1 | 11,920 |
| BCC 32 | *Cx. annulirostris* | Bracken R | 5/1/2022 | 200 | RRV | RRV | 40,836,524 | 1,474,766 | 3.61 | 9,279 | 11,911 | MH987781.1 | 11,920 |
| BCC 45 | *Ae. procax* | Corinda | 17/1/2022 | 160 | RRV | RRV | 40,000,705 | 881,589 | 2.20 | 5,547 | 11,911 | MH987781.1 | 11,920 |
| BCC 48 | *Ae. procax* | Corinda | 7/2/2022 | 117 | RRV | RRV | 40,922,299 | 390,761 | 0.95 | 2,459 | 11,912 | MH987781.1 | 11,920 |
| BCC 63 | *Cx. annulirostris* | Indooroopilly | 14/2/2022 | 135 | RRV | RRV | 35,404,670 | 275,631 | 0.78 | 1,734 | 11,912 | MH987781.1 | 11,920 |
| BCC 70 | *Cx. annulirostris* | Banyo | 21/2/2022 | 200 | RRV | RRV | 39,348,668 | 963,678 | 2.45 | 6,063 | 11,911 | MH987781.1 | 11,920 |
| BCC 83 | *Ae. vigilax* | Banyo | 11/4/2022 | 104 | RRV | RRV | 42,975,529 | 625,696 | 1.46 | 3,937 | 11,911 | MH987781.1 | 11,920 |
| QH 138 | *Cx. annulirostris* | Bracken R | 4/2/2022 | 200 | RRV | RRV | 46,412,925 | 622,180 | 1.34 | 3,915 | 11,912 | MH987781.1 | 11,920 |
| QH 329 | *Cx. annulirostris* | Indooroopilly | 9/3/2022 | 200 | RRV | RRV | 41,290,576 | 60,550 | 0.15 | 381 | 11,908 | MH987781.1 | 11,920 |
| BCC 380 | *Cx. orbostiensis* | Hemmant | 17/5/2021 | 1 | RRV | no sequence obtained | 40,110,924 | - | - | - | - | - | - |
| BCC 317 | *Cx. sitiens* | Indooroopilly | 14/2/2022 | 3 | STRV & RRV | no sequence obtained | 35,639,262 | - | - | - | - | - | - |
| QH 160 | *Ve. funerea* | Bracken R | 4/2/2022 | 58 | BFV | BFV | 40,821,131 | 2,974 | 0.01 | 19 | 11,549 | MN064696.1 | 11,574 |
| BCC 71 | *Cx. annulirostris* | Banyo | 21/2/2022 | 200 | BFV | BFV | 39,117,880 | 4,476 | 0.01 | 29 | 11,554 | MN064696.1 | 11,574 |
| BCC 22 | *Ae. vigilax* | Bracken R | 24/5/2021 | 125 | BFV | BFV | 41,718,056 | 49,426 | 0.12 | 320 | 11,555 | MN064696.1 | 11,574 |
|  |  |  |  |  |  | STRV | 41,718,056 | 3,657 | 0.01 | 25 | 10,830 | MZ358850.1 | 10,846 |
| BCC 76 | *Ae. procax* | Banyo | 11/4/2022 | 200 | STRV | STRV | 40,014,774 | 38,728 | 0.10 | 268 | 10,846 | MZ358850.1 | 10,846 |
| QH 158 | *Ae. procax* | Bracken R | 4/2/2022 | 202 | STRV | STRV | 43,652,133 | 2,153 | 0.005 | 15 | 10,803 | MZ358850.1 | 10,846 |
| QH 226 | *Ae. procax* | Banyo | 3/2/2022 | 132 | STRV | STRV | 39,253,971 | 2,326 | 0.10 | 16 | 10,846 | MZ358850.1 | 10,846 |
| QH 295 | *Ae. procax* | Corinda | 17/2/2022 | 190 | STRV | STRV | 32,878,765 | 15,082 | 0.05 | 104 | 10,846 | MZ358850.1 | 10,846 |
| BCC 67 | *Cx. annulirostris* | Banyo | 21/2/2022 | 200 | SINV | SINV | 34,843,064 | 6,429 | 0.02 | 42 | 11,604 | OP950209.1 | 11,611 |

Depth of coverage = Number of mapped reads × 75 (read length)/reference genome length. QTD: quantity.
BCC (collected by Brisbane City Council) = samples with previous detection of RRV in FTA cards
QH (collected by Queensland Health) = samples collected after detection of RRV in FTA cards

For the 20 mosquito pools further investigated by metatranscriptomic analysis, a mean of 39,622,773 paired reads was generated per sample (range: 32,878,765– 46,412,925). We generated 18 novel full-length sequences, with some samples containing up to two different viruses (**Table 2**). Full-length genomes displayed an average coverage depth of 2,248 reads (range: 15–9,279), with genome size ranging from 10.803 to 11,913 nucleotides (**Table 2**).

### 3.3. Phylogenetic analyses

The 18 complete genomes generated in this study (9 of RRV; 3 of BFV; 1 of SINV-like; and 5 of STRV) were used to reconstruct the phylogenetic relationship of these viruses in comparison with the genomic sequences available in public databases. Overall, phylogenetic analysis suggested continuous circulation of all arboviruses over the last 90 years in Australia. The lineages that are currently circulating in the country are RRV_G4B, SINV_G3C, STRV_G2B, and BFV_G3B, which all emerged within the last 26 to 11 years (**Figures 2 and S2**). Phylogeographic analysis revealed long-distance dispersal of all lineages within Australia (**Table S6; Figure S1**).

**Figure 2.**
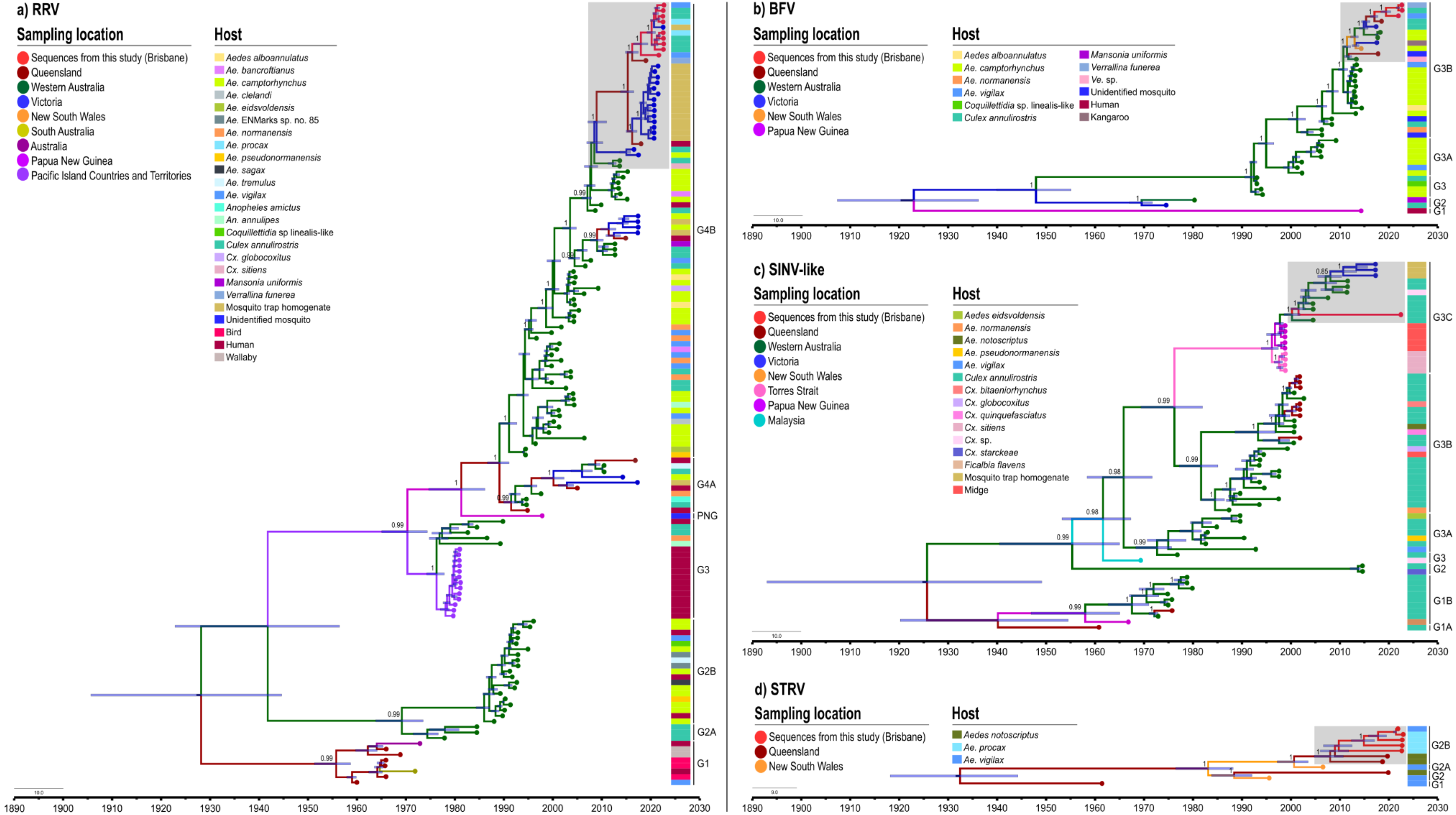
Maximum clade credibility tree based on the phylogenetic analysis of complete polyprotein sequences of (**a**) Ross River virus (RRV), (**b**) Barmah Forest virus (BFV), (**c**) Sindbis-like virus (SINV-like) and (**d**) Stratford virus (STRV). Nodes are coloured based on their geographical origin, as indicated in the map of Australia showed previously. Coloured squares represent host origin. Grey shaded squares represent clades circulating in the study area. Posterior probability values of >0.80 are presented above main branches.

Except for STRV, all the other viruses investigated showed a clear ladder-like tree structure (**Figure 2**) which is characteristic of RNA virus evolution under strong immunological constraints (from both mosquitoes and vertebrate hosts) leading to continuous lineage turnover overtime (Makau et al., 2022).

#### 3.3.1. Ross River virus

A total of 141 RRV whole genome sequences from publicly available data bases were analysed, including the nine new genomes generated here. The virus sequences originated from Papua New Guinea (PNG), the Pacific Island Countries and Territories (PICTs) and different states of Australia, including New South Wales (NSW), Queensland (QLD), South Australia (SA), Victoria (VIC) and Western Australia (WA) (**Figure S3**). A maximum likelihood (ML) phylogeny was re-constructed showing that the sequences obtained in this study belonged to the G4 genotype, which encompasses all contemporary (1994-2022) RRV isolates (Michie et al., 2021). Our samples belonged to the G4B lineage, which includes mosquito-derived strains from QLD, VIC and WA, and human-derived strains from QLD and WA (**Figure S4**).

The time to the most recent common ancestor (TMRCA) of all RRV genomes was estimated to be around 1927 (95% highest posterior density [HPD] = 1908 to 1943) and the current subclade of G4B has been circulating since ∼ 2007 (95% HPD = 2005 to 2008) (**Figure 2a**).

#### 3.3.2. Barmah Forest virus

A total of 39 BFV whole genome sequences were analysed. The strains were sampled from PNG and from the Australian states of NSW, QLD, VIC and WA, between 1974–2018, including our three new genomes from 2021–2022 (**Figure S5**). The sequences obtained in this study belong to G3 genotype, G3B lineage, to which all contemporary BFV strains in Australia belong (Michie et al., 2020). This lineage consists of mosquito-derived strains from QLD, VIC, and WA, and a kangaroo-derived strain from NSW (**Figure S6)**. The TMRCA of all BFV genomes was estimated to occur around 1922 (95% HPD = 1907 to 1936) and the current subclade of G3B emerged around 2011 (95% HPD = 2010 to 2012) (**Figure 2b**).

#### 3.3.3. Sindbis-like virus

All available whole and draft genome (>10 kb) sequences of Sindbis and Sindbis-like viruses were analysed. The final alignment contained 161 sequences from Australasia, Africa and Europe, described between 1953–2022 (**Figure S7**). Phylogenetic analyses revealed four major genetic groups with more than 23% nucleotide sequence divergence and 8% amino acid sequence divergence between these groups, across their entire genomes. According to the International Committee on Taxonomy of Viruses this meets the criteria for demarcating Alphavirus species (Powers et al., 2009). These new species have been described by Michie et al., 2023.

Our sample belongs to the most recent lineage of SINV-like present in Australia, known as G3C (**Figure S8**). This lineage consists of mosquito-derived strains, mostly from *Cx. annulirostris*, from QLD, VIC, and WA, and midge-derived strains from PNG (**Figure S9)**. The TMRCA of all SINV-like lineages of Australia was estimated to be approximately 1924 (95% highest posterior density (95% HPD = 1886 to 1949). The emergence of G3C dates to approximately 1996 (95% HPD = 1994 to 1998), and the current subclade has been circulating since 2000 (95% HPD = 1998 to 2002) (**Figure 2c**).

#### 3.3.4. Stratford virus

A total of 11 STRV whole genome sequences sampled from NSW and QLD, between 1961–2019, including the five new genomes generated in this study (from 2021–2022) were analysed (**Figure S10**). Despite the analysis being limited to only eleven STRV isolates, phylogenetic analyses revealed genetic diversity in mosquito-derived samples from QLD and NSW. The TMRCA of STRV was estimated around 1932 (95% HPD = 1918 to 1944). The current subclade, G2B, has been circulating since 2007 (95% HPD = 1997 to 2003) (**Figure 2d**).

## 4. Discussion

Arboviruses pose a significant threat globally. There are more than 500 known arboviruses of which approximately 100 are pathogenic to humans (Artsob et al., 2023). In Australia, at least thirteen arboviruses have been associated with human disease. Some of the endemic zoonotic arboviruses, such as BFV and RRV, are maintained in complex, poorly understood transmission cycles, involving a broad range of mosquito species and potential vertebrate hosts (Mackenzie et al., 1994; Ong et al., 2021). As a result, transmission pathways may differ geographically and temporally, leading to variable human transmission risks. Here, we successfully generated 18 novel full-length genome sequences, the phylogenetic analysis of which increased our current understanding of Australian arbovirus evolution. These sequences and their analyses highlight major knowledge gaps in arbovirus phylogeny in Australia and unveil an evolutionary history marked by continuous transmission, co-circulation, and steady lineage modification over time.

The arboviruses found in our study are widespread within Australia. While their exact geographic origins and route of spread could not be determined due to limited whole-genome sequence information, phylogeographic analysis indicated long-distance movement of all lineages within the country probably facilitated by the movement of viraemic hosts (including mosquitoes, birds, humans and other mammals). Isolates from widely separated locations, shared similar genomic sequences and lineages, and suggested evolution over a large temporal timeframe (100 years) (**Figure 2 and S1**). Genomes sequenced by this project were recovered from *Cx. annulirostris*, *Ae. vigilax*, *Ae. procax* and *Ve. funerea*. All are competent vectors of at least a subset of the viruses isolated (Harley et al., 2001) and might play key roles in establishing local transmission across parts their ranges. However, further studies, and many more genomes are needed to characterise the role of individual reservoirs and vector species in maintaining the transmission of specific arbovirus lineages in different habitats.

The Australian alphaviruses phylogenies exhibit a ladder-like structure, where a single dominant lineage links viruses sampled from various time points. This pattern, observed in other viruses (e.g., influenza virus), is consistent with strongly immune-driven evolution (Grenfell et al., 2004; Volz et al., 2013). The phylogenetic trees of arboviruses are inherently complex due to the many components of transmission. For instance, DENV faces selection pressures from the immune systems of both its humans and mosquito hosts, resulting in novel virus variants (Stica et al., 2022; Wash and Soria, 2015). This complexity is particularly pronounced for zoonotic arboviruses because they have multiple vectors and reservoirs. The long-term transmission of these viruses in Australia, their high seroprevalence in vertebrate host species (Vieira et al., 2024) and the shape of their phylogenetic trees supports the hypothesis of a strong immunological barrier driving lineage evolution (antigenic drift) and turnover, at least for BFV, RRV and SINV.

While phylogenetic trees serve as useful indicators of epidemiological, immunological, and evolutionary processes affecting viral genetic variation, ladder-like trees (e.g., **Figure 2 a–c**), can indicate the presence of directional selection, but can also reflect the sequential genetic bottlenecks associated with rapid spatial spread (e.g., rabies virus; (Streicker et al., 2010)). To further understand phylogenetic tree shapes and test these hypotheses, more genomic surveillance is required. Exploring these factors would provide valuable insights into arboviral transmission dynamics.

Similar divergence dates were estimated for all the arboviruses identified in this study, with genotypes diverging around 1922-1932. Coincidentally, this timeframe aligns with the first reports of "an unusual epidemic" in Australia, with the syndrome of polyarthralgia and rash (Nimmo, 1928). The causal agent of the disease was later suggested to be RRV (Doherty et al., 1971, 1964), but could have been caused by any “indigenous pathogen” (Jacups et al., 2008). While some major ecological event may have led to the emergence of arboviruses at this time, we also note that 100 years may simply represent the limits of our methodology: rapid viral evolution allows us to recover past estimates of viral emergence and divergence but once signal saturation is reached it limits accurate estimates of ancient viral emergence at the distant past (Aiewsakun and Katzourakis, 2015).

### 4.1. Virus screening and phylogenetic analyses

Ross River virus is responsible for the highest number of human arbovirus infection notifications across every state and territory of Australia (Jansen et al., 2019) (**Figure S11**). The virus was first isolated in 1959, from a pool of *Ae. vigilax* mosquitoes collected from Queensland (Doherty et al., 1963) and from humans in 1972 in the same state (Doherty et al., 1972). Over 40 species of mosquito and 20 vertebrate hosts have been associated with RRV transmission (Claflin and Webb, 2015; Stephenson et al., 2019). Sixty-five years after its isolation, Australia remains no closer to understanding transmission, predicting human spillover, or implementing a surveillance program that could enhance our understanding. In this study, five mosquito species found to harbour RRV, four of which have been previously identified as competent RRV vectors in laboratory studies (reviewed in Russell, 2002). Of these, *Ae. vigilax* and *Cx. annulirostris*, yield the most field detections and vector competence studies have implicated these as vector species, along with *Ae. procax* and *Cx. sitiens.* We also detected RRV by PCR in *Cx. orbostiensis*, but no laboratory studies on the vector competence of this species have been conducted.

Phylogenetic reconstructions were made using RRV sequences from 20 mosquito species, birds, humans, and wallabies. All belonged to the G4 genotype, G4B lineage. As described by Michie et al. (2021), all strains collected in Australia since 1996 belong to G4 genotype, indicating that it is the contemporary and dominant genotype in circulation in the country. The last Australian detection of G4A occurred in 2016 (VIC and QLD) and all recently sampled strains, including sequences from QLD (2016–2018), VIC (2016–2017), WA (2008– 2013), and this study (2021–2022), were classified as G4B. The G4B lineage have been found in samples from humans and 15 mosquito species, which underscores the complexity of RRV’s vector range but there are no sequences from non-human vertebrate hosts, highlighting a key gap in knowledge regarding reservoir incrimination.

Barmah Forest virus is responsible for the second highest number of human arbovirus notifications in Australia (Knope et al., 2019) (**Figure S12**). This alphavirus was first isolated in 1974 from *Cx. annulirostris* mosquitoes collected from Victoria (VIC) (Marshall et al., 1982) and concurrently from mosquitoes trapped in Queensland (Doherty et al., 1979). The first case of BFV infection in humans was not reported until 1986 (Boughton et al., 1988). The virus has been isolated from several wild-caught mosquito species, including those found positive for the virus in this study: *Cx. annulirostris, Ae. vigilax* and *Ve. funerea* (Jacups et al., 2008; Jeffery et al., 2006). The sequences obtained in this study belong to the G3 genotype, G3B lineage. G3 is the contemporary genotype circulating in Australia (Michie et al. (2020)) and the G3B is widely distributed. The number of sequences of the G3B lineage is limited but originate from a kangaroo and six mosquito species. More genomic information is needed to understand the potential transmission dynamics of this virus.

Sindbis virus (here called Sindbis-like virus due to its divergence from the Europe-African virus group (Michie et al., 2023)) is an alphavirus first isolated in Australia in 1960 from *Cx. annulirostris* mosquitoes (Doherty et al., 1963a). Approximately 65% of SINV-like genomes investigated in Australia have been isolated from this mosquito, including the strains found in this study. It is one of the most commonly isolated arboviruses in Australian mosquitoes (Ong et al., 2021). The genotype detected in this study, G3C, circulates in WA and has also been detected in mosquitoes from VIC in 2016 (Batovska et al., 2022). Although SINV-like virus appears to have an association with mild infection in humans (Doherty, 1973; Doherty et al., 1969; Guard et al., 1982), the resulting health implications remain unclear, and its vertebrate reservoirs are unknown. No outbreaks of SINV-like virus have been reported in Australia, despite clear evidence that this virus has been circulating in the country for over a century. It is not notifiable in the country and there is no clinical or widely available laboratory diagnostic.

Stratford virus was the only flavivirus found in our study. It is a member of the Kokobera virus subgroup and was first isolated in 1961 from *Ae. vigilax* mosquitoes collected in north Queensland (Doherty et al., 1963a). It is often detected in *Aedes* species, but information regarding vertebrate reservoirs is not available (Toi et al., 2017). Based on serological evidence, sporadic human infections with STRV have been documented in asymptomatic individuals in NSW (Hawkes et al., 1985), as well as in symptomatic patients in NSW and QLD with symptoms including fever, joint pain and lethargy (Johansen et al., 2005; Phillips et al., 1993; Pyke et al., 2021). Notifications designated as “unspecified-flavivirus” infections, which includes STRV, are frequently submitted to the Australian Government National Notifiable Disease Surveillance System. These cases are confirmed through serology, and about 30 unspecified-flavivirus infections are reported annually reaching its peak (116 cases) in 2016 (Department of Health and Aged Care, 2023) (**Figure S13**). Between 2017 and 2020, a total of 49 suspected-flavivirus infections in QLD were STRV IgM positive (Pyke et al., 2021).

Although limited to a comparison of eleven available contemporary STRV full sequences, phylogenetic analysis demonstrated genetic diversity among the respective QLD and NSW isolate groups. Together with our six new isolations in Brisbane, these findings support evidence of ongoing circulation of STRV. The geographical distribution of the virus is likely to be broader than currently described and potential vertebrate reservoirs remain unknown. Further genomic surveillance would improve our understanding of STRV circulation.

### 4.2. The role of xeno-monitoring coupled with metatranscriptomics in disease surveillance

Xeno-surveillance of mosquito-borne pathogens in Australia has been underway for many decades, but state programs differ in magnitude and by methodological sensitivity. Routine screening of mosquitoes (by trap homogenate or by FTA card) only occurs for a subset of “notifiable” Australian endemic viruses: BFV, RRV, JEV, WNVKUN, and MVEV (Department of Health and Aged Care, 2023). These are PCR tests for virus fragments, often in undifferentiated mosquito pools (i.e., no data on species associations) and, routinely, detections are not investigated by genome sequencing. For instance, during the 2022-2023 surveillance season, a total of 179 arbovirus detections in mosquitos were recorded in the states of NSW, VIC and South Australia (SA) (Government of South Australia, 2023; NSW Government, 2023; Victoria State Government, 2023) (**Table S7**), but no genomic data is available. No public reports on xeno-surveillance programs from other states are available. Non-notifiable arboviruses circulating in Australia, including SINV-like and STRV, are not routinely tested for using xeno-diagnostic surveillance, possibly because, to date, their pathogenicity in humans appears mild and is poorly described. In the absence of comprehensive surveillance programs, current xeno-surveillance programs tend simply to identify the presence of virus in complex environments that are already known to be endemic. The key transmission pathways and distributions of specific genetic lineages remains unknown (Gyawali et al., 2019). Nonetheless, there is increasing awareness of the crucial nature of molecular xeno-monitoring in response to the emergence and re-emergence of neglected arboviruses (Hill et al., 2023; Laiton-Donato et al., 2023). This is of particular significance on the Australian continent where a large number of ecosystems, at huge geographic scales, are under constant environmental disturbance potentially favoring arbovirus emergence and dispersal through dynamic changes in mosquito vector populations, reservoir distribution, and human proximity to those reservoirs and vectors. Designing a sustainable but informative surveillance system is a tremendous challenge.

The COVID-19 pandemic has highlighted the utility of genomic surveillance in tracking and monitoring the spread of viruses, detecting new variants, and informing public health interventions (Hill et al., 2023; Zeghbib et al., 2023). However, adapting routine sequencing into public health investigations will require additional programmatic investment in expertise and resources. We demonstrate that pan-alphavirus and pan-flavivirus screens can be used to test large but differentiated mosquito pools for arboviruses. Further investigations on genomes are then conducted on positive samples. Sequencing the “known” positives reduces the costs associated with NGS and enhances the efficiency of the surveillance network, yielding more informative results. In the future, multi-locus DNA metabarcoding approaches, similar to those demonstrated in conservation science (Arulandhu et al., 2017) might increase efficiencies further by initial screening that include the identification of vectors, vertebrate signals and pathogens from the same trap collection (Arulandhu et al., 2017).

Whole genome sequencing will improve existing information available for public health. It provides significant phylogenetic resolution that enables the reconstruction of local transmission chains, determination of the geographical origin of virus emergence and transmission, tracking of virus mutations, and identification of viral strains with modified phenotype (Pollett et al., 2020). Viruses such as RRV, BFV, JEV and MVEV, are primarily maintained by active infection in animal reservoirs, which may, especially in the case of RRV, include humans (**Figure S14 and S15**) (Mackenzie et al., 1994). Although these viruses have encephalitogenic and arthritogenic symptoms of public health importance (see Mcguinness et al., 2023 for JEV and MVEV review, and Ong et al., 2021 for RRV review), there is limited whole genome information for each virus, especially for MVEV and JEV in Australia (**Figure S16 and S17** respectively). Xeno-monitoring can uncover ecological factors that contribute to outbreaks, including cryptic transmission of viral lineages and transmission patterns (Cameron and Ramesh, 2021). Ultimately, effective genomic xeno-monitoring programs have the potential to generate information for building risk models that can target public health messaging, vector control, and vaccination to prevent and mitigate the impact of arboviruses (Grubaugh et al., 2019a).

## 5. Conclusion

The simultaneous circulation of multiple arboviruses and a limited understanding of temporal transmission dynamics indicate that improved, well-designed surveillance programs of arboviruses that include genomic surveillance, are needed. Curation of longitudinal, consistent data sets that are shared with the scientific community are invaluable (Cameron and Ramesh, 2021; Laiton-Donato et al., 2023). To gain a comprehensive understanding of Australian arboviruses phylogeography and movement patterns across the country, routine sampling and sequencing of viruses from the different states and territories, over time is necessary. This analysis would enable the identification of potential foci for viral diversity generation and the detailed source and sink hubs for transmission within Australia and the region. Finally, integrating sequencing approaches into arbovirus surveillance strategies around Australia may provide high-resolution data for researchers and public health agencies to better understand the emergence and evolution of viruses, ultimately enhancing preparedness and response strategies to mitigate the impact of future outbreaks.

## Data availability

Genome sequences obtained in this study have been deposited in GenBank under accession numbers PP496979 to PP496996 and the raw NGS reads are available the NCBI Sequence Read Archive (SRA) database under BioProject PRJNA1077787. Metadata used and generated in this study is available at https://github.com/carlavieira1/Brisbane-arboviruses---metadata.

## Acknowledgements

The authors acknowledge the contribution of the Communicable Disease Branch of Queensland Health who support Queensland’s arbovirus surveillance program by screening FTA cards. C.J.S.P.V. was supported by a QIMRB International PhD Scholarship, QUT Tuition Fee Sponsorship, and funds from the Mosquito Control Laboratory, QIMRB.

## Author contributions

G.J.D. devised the project. G.L.W., A.F.v.d.H., F.D.F. and G.J.D. supervised the project. C.J.S.P.V., M.B.O., M.A.S., D.S. and J.M.D. coordinated field activities. C.J.S.P.V. identified, pooled, and processed mosquito samples. C.J.S.P.V. and M.G. prepared library. C.J.S.P.V. collated and analysed the data, drafted the manuscript and designed the figures. M.B.O., G.L.W., A.F.v.d.H., F.D.F., and G.J.D. contributed to the editing of the final manuscript. All authors discussed the results and approved the version submitted for publication.

## References

Aiewsakun, P., Katzourakis, A., 2015. Endogenous viruses: Connecting recent and ancient viral evolution. Virology 479–480, 26–37.

Artsob, H., Lindsay, R., Drebot, M., 2023. Arboviruses. In: Reference Module in Biomedical Sciences. Elsevier.

Arulandhu, A.J., Staats, M., Hagelaar, R., Voorhuijzen, M.M., Prins, T.W., Scholtens, I., Costessi, A., Duijsings, D., Rechenmann, F., Gaspar, F.B., Crespo, M.T.B., Holst-jensen, A., Birck, M., Burns, M., Haynes, E., Hochegger, R., Klingl, A., Lundberg, L., Natale, C., Niekamp, H., Perri, E., Barbante, A., Rosec, J., Seyfarth, R., Sovova, T., Moorleghem, C. Van, Ruth, S. van, Peelen, T., Kok, E., 2017. Development and validation of a multi-locus DNA metabarcoding method to identify endangered species in complex samples. Gigascience 6, 1–18.

Australian Bureau of Statistics, 2022. 2021 Census QuickStats [WWW Document]. Aust. Bur. Stat. URL https://www.abs.gov.au/census/find-census-data/quickstats/2021/3GBRI (accessed 4.4.23).

Batovska, J., Lynch, S.E., Cogan, N.O.I., Brown, K., Darbro, J.M., Kho, E.A., Blacket, M.J., 2018. Effective mosquito and arbovirus surveillance using metabarcoding. Mol. Ecol. Resour. 18, 32–40.

Batovska, J., Mee, P.T., Lynch, S.E., Sawbridge, T.I., Rodoni, B.C., 2019. Sensitivity and specificity of metatranscriptomics as an arbovirus surveillance tool. Sci. Rep. 9, 19398.

Batovska, J., Mee, P.T., Sawbridge, T.I., Rodoni, B.C., Lynch, S.E., 2022. Enhanced Arbovirus Surveillance with High-Throughput Metatranscriptomic Processing of Field-Collected Mosquitoes. Viruses 14, 2759.

BoM, B. of M., 2023. Climate statistics for Australian locations. Monthly climate statistics. [WWW Document]. Aust. Gov. Bur. Meteorol. URL http://www.bom.gov.au/climate/averages/tables/cw_040913.shtml (accessed 4.4.23).

Boughton, C.R., Hawkes, R.A., Nairn, H.M., 1988. Illness caused by a Barmah Forest-like virus in New South Wales. Med. J. Aust. 148, 146–147.

Cameron, M.M., Ramesh, A., 2021. The use of molecular xenomonitoring for surveillance of mosquito-borne diseases. Philos. Trans. R. Soc. B Biol. Sci. 376, 20190816.

CDC, C. for D.C. and P., 2023. Arbovirus Catalog [WWW Document]. Centers Dis. Control Prev. URL https://wwwn.cdc.gov/arbocat/ (accessed 3.28.23).

Claflin, S.B., Webb, C.E., 2015. Ross River Virus: Many Vectors and Unusual Hosts Make for an Unpredictable Pathogen. PLOS Pathog. 11, e1005070.

Department of Health and Aged Care, 2023. Australian Government. National Communicable Disease Surveillance Dashboard [WWW Document]. URL https://www.health.gov.au/our-work/nndss (accessed 3.16.23).

Dezordi, F.Z., Neto, A.M. da S., Campos, T. de L., Jeronimo, P.M.C., Aksenen, C.F., Almeida, S.P., Wallau, G.L., 2022. ViralFlow: A Versatile Automated Workflow for SARS-CoV-2 Genome Assembly, Lineage Assignment, Mutations and Intrahost Variant Detection. Viruses 14, 217.

Doherty, R.L., 1973. Surveys of haemagglutination-inhibiting antibody to arboviruses in aborigines and other population groups in Northern and Eastern Australia, 1966-1971. Trans. R. Soc. Trop. Med. Hyg. 67, 197–205.

Doherty, R.L., Barrett, E.J., Gorman, B.M., Whitehead, E.H., 1971. Epidemic polyarthritis in Eastern Australia, 1959-1971. Med. J. Aust. 1, 5–8.

Doherty, R.L., Bodey, A.S., Carew, J.S., 1969. Sindbis virus infection in Australia. Med. J. Aust. 2, 1016–1017.

Doherty, R.L., Carley, J.G., Best, J.C., 1972. Isolation of Ross River virus from man. Med. J. Aust. 1, 1083–1084.

Doherty, R.L., Carley, J.G., Kay, B.H., Filippich, C., Marks, E.N., Frazier, C.L., 1979. Isolation of virus strains from mosquitoes collected in Queensland, 1972-1976. Aust. J. Exp. Biol. Med. Sci. 57, 509–520.

Doherty, R.L., Carley, J.G., Mackerras, M.J., Marks, E.N., 1963a. Studies of arthropod-borne virus infections in Queensland. III. Isolation and characterization of virus strains from wild-caught mosquitoes in North Queensland. Aust. J. Exp. Biol. Med. Sci. 41, 17–39.

Doherty, R.L., Gorman, B.M., Whitehead, R.H., Carley, J.G., 1964. Studies of epidemic polyarthritis: the significance of three group A arboviruses isolated from mosquitoes in Queensland. Australas. Ann. Med. 13, 322–327.

Doherty, R.L., Whitehead, R.H., Gorman, B.M., O’Gower, A.K., 1963b. The isolation of a third group A arbovirus in Australia, with preliminary observations on its relationship to epidemic polyarthritis. Aust. J. Sci. 26, 183–184.

Faria, N.R., Quick, J., Claro, I.M., Thézé, J., Jesus, J.G. de, Giovanetti, M., Kraemer, M.U.G., Hill, S.C., Black, A., Costa, A.C. da, Franco, L.C., Silva, S.P., Wu, C.-H., Raghwani, J., Cauchemez, S., Plessis, L. du, Verotti, M.P., Oliveira, W.K. de, Carmo, E.H., Coelho, G.E., Santelli, A.C.F.S., Vinhal, L.C., Henriques, C.M., Simpson, J.T., Loose, M., Andersen, K.G., Grubaugh, N.D., Somasekar, S., Chiu, C.Y., Muñoz-Medina, J.E., Gonzalez-Bonilla, C.R., Arias, C.F., Lewis-Ximenez, L.L., Baylis, S.A., Chieppe, A.O., Aguiar, S.F., Fernandes, C.A., Lemos, P.S., Nascimento, B.L.S., Monteiro, H.A.O., Siqueira, I.C., Queiroz, M.G. de, Souza, T.R. de, Bezerra, J.F., Lemos, M.R., Pereira, G.F., Loudal, D., Moura, L.C., Dhalia, R., França, R.F., Magalhães, T., Marques-Jr, E.T., Jaenisch, T., Wallau, G.L., Lima, M.C. de, Nascimento, V., Cerqueira, E.M. de, Lima, M.M. de, D. L. Mascarenhas, J. P. Moura Neto, A. S. Levin, T. R. Tozetto-Mendoza, S. N. Fonseca, M. C. Mendes-Correa, F. P. Milagres, A. Segurado, E. C. Holmes, A. Rambaut, T. Bedford, M. R. T. Nunes, E. C. Sabino, L. C. J. Alcantara, N. J. Loman & O. G. PybusN. R, E.C.H., Rambaut, A., Bedford, T., Nunes, M.R.T., Sabino, E.C., Alcantara, L.C.J., Loman, N.J., Pybus, O.G., 2017. Establishment and cryptic transmission of Zika virus in Brazil and the Americas. Nature 546, 406–410.

Furuya-kanamori, L., Gyawali, N., Mills, D.J., Hugo, L.E., Devine, G.J., Lau, C.L., 2022. The Emergence of Japanese Encephalitis in Australia and the Implications for a Vaccination Strategy. Trop. Med. Infect. Dis. 7, 85.

Government of South Australia, S.H., 2023. The South Australian arbovirus and mosquito monitoring report.

Grenfell, B.T., Pybus, O.G., Gog, J.R., Wood, J.L.N., Daly, J.M., Mumford, J.A., Holmes, E.C., 2004. Unifying the Epidemiological and Evolutionary Dynamics of Pathogens. Science (80-.). 303, 327–332.

Grubaugh, N.D., Ladner, J.T., Lemey, P., Pybus, O.G., Rambaut, A., Holmes, E.C., Andersen, K.G., 2019a. Tracking virus outbreaks in the twenty-first century. Nat. Microbiol. 4, 10–19.

Grubaugh, N.D., Saraf, S., Gangavarapu, K., Watts, A., Tan, A.L., Oidtman, R.J., Ladner, J.T., Oliveira, G., Matteson, N.L., Kraemer, M.U.G., Vogels, C.B.F., Hentoff, A., Bhatia, D., Stanek, D., Scott, B., Landis, V., Stryker, I., Cone, M.R., Kopp, E.W., Cannons, A.C., Heberlein-Larson, L., White, S., Gillis, L.D., Ricciardi, M.J., Kwal, J., Lichtenberger, P.K., Magnani, D.M., Watkins, D.I., Palacios, G., Hamer, D.H., Gardner, L.M., Perkins, T.A., Baele, G., Khan, K., Morrison, A., Isern, S., Michael, S.F., Andersen, K.G., 2019b. Travel Surveillance and Genomics Uncover a Hidden Zika Outbreak during the Waning Epidemic. Cell 178, 1057–1071.e11.

Guard, R.W., Mcauliffe, M.J., Stallman, N.D., Bramston, B.A., 1982. Haemorrhagic manifestations with sindbis infection. Case report. Pathology 14, 89–90.

Gubler, D.J., 1998. Dengue and Dengue Hemorrhagic Fever. Clin. Microbiol. Rev. 11, 480– 496.

Guindon, S., Dufayard, J.-F., Lefort, V., Anisimova, M., Hordijk, W., Gascuel, O., 2010. New Algorithms and Methods to Estimate Maximum-Likelihood Phylogenies: Assessing the Performance of PhyML 3.0. Syst. Biol. 59, 307–321.

Gyawali, N., Taylor-Robinson, A.W., Bradbury, R.S., Pederick, W., Faddy, H.M., Aaskov, J.G., 2019. Neglected Australian Arboviruses Associated With Undifferentiated Febrile Illnesses. Front. Microbiol. 10, 2818.

Hall-Mendelin, S., Ritchie, S.A., Johansen, C.A., Zborowski, P., Cortis, G., Dandridge, S., Hall, R.A., Van Den Hurk, A.F., 2010. Exploiting mosquito sugar feeding to detect mosquito-borne pathogens. Proc. Natl. Acad. Sci. U. S. A. 107, 11255–11259.

Harley, D., Sleigh, A., Ritchie, S., 2001. Ross river virus transmission, infection, and disease: A cross-disciplinary review. Clin. Microbiol. Rev. 14, 909–932.

Hawkes, R.A., Boughton, C.R., Naim, H.M., Wild, J., Chapman, B., 1985. Arbovirus infections of humans in New South Wales. Seroepidemiology of the flavivirus group of togaviruses. Med. J. Aust. 143, 555–561.

Hermanns, K., Zirkel, F., Kopp, A., Marklewitz, M., Rwego, I.B., Estrada, A., Gillespie, T.R., Drosten, C., Junglen, S., 2017. Discovery of a novel alphavirus related to Eilat virus. J. Gen. Virol. 98, 43–49.

Hill, V., Githinji, G., Vogels, C.B.F., Bento, A.I., Chaguza, C., Carrington, C.V.F., Grubaugh, N.D., 2023. Toward a global virus genomic surveillance network. Cell Host Microbe 31, 861–873.

Huang, Y.S., Higgs, S., Vanlandingham, D.L., 2019. Emergence and re-emergence of mosquito-borne arboviruses. Curr. Opin. Virol. 34, 104–109.

Jacups, S.P., Whelan, P.I., Currie, B.J., 2008. Ross River virus and Barmah Forest virus infections: a review of history, ecology, and predictive models, with implications for tropical northern Australia. Vector borne zoonotic Dis. 8, 283–297.

Jansen, C.C., Shivas, M.A., May, F.J., Pyke, A.T., Onn, M.B., Lodo, K., Hall-Mendelin, S., McMahon, J.L., Montgomery, B.L., Darbro, J.M., Doggett, S.L., Van Den Hurk, A.F., 2019. Epidemiologic, entomologic, and virologic factors of the 2014-15 Ross River virus outbreak, Queensland, Australia. Emerg. Infect. Dis. 25, 2243–2252.

Jeffery, J.A.L., Kay, B.H., Ryan, P.A., 2006. Role of Verrallina funerea (Diptera: Culicidae) in Transmission of Barmah Forest Virus and Ross River Virus in Coastal Areas of Eastern Australia. J. Med. Entomol. 43, 1239–1247.

Johansen, C.A., Maley, F.M., Broom, A.K., 2005. Isolation of Stratford Virus from Mosquitoes Collected in the Southwest of Western Australia. Arbovirus Res. Aust. 9, 164–166.

Katoh, K., Rozewicki, J., Yamada, K.D., 2019. MAFFT online service: Multiple sequence alignment, interactive sequence choice and visualization. Brief. Bioinform. 20, 1160– 1166.

Kizu, J.G., Graham, M., Grant, R., Mccallum, F., Mcpherson, B., Auliff, A., Kaminiel, P., Liu, W., 2023. Prevalence of Barmah Forest Virus, Chikungunya Virus and Ross River Virus Antibodies among Papua New Guinea Military Personnel before 2019 †. Viruses 15, 394.

Knope, K., Doggett, S.L., Jansen, C.C., Johansen, C.A., Kurucz, N., Feldman, R., Lynch, S.E., Hobby, M.P., Sly, A., Jardine, A., Bennett, S., Currie, B.J., Toms, C., 2019. Arboviral diseases and malaria in Australia, 2014–15: Annual report of the National Arbovirus and Malaria Advisory Committee. Commun. Dis. Intell. 43.

Laiton-Donato, K., Guzmán-Cardozo, C., Peláez-Carvajal, D., Ajami, N.J., Navas, M.C., Parra-Henao, G., Usme-Ciro, J.A., 2023. Evolution and emergence of mosquito-borne viruses of medical importance: Towards a routine metagenomic surveillance approach. J. Trop. Ecol. 39, e13.

Langat, S.K., Eyase, F.L., Berry, I.M., Nyunja, A., Bulimo, W., Owaka, S., Ofula, V., Limbaso, S., Lutomiah, J., Jarman, R., Distelhorst, J., Sang, R.C., 2020. Origin and evolution of dengue virus type 2 causing outbreaks in Kenya: Evidence of circulation of two cosmopolitan genotype lineages. Virus Evol. 6, veaa026.

Larsson, A., 2014. AliView: A fast and lightweight alignment viewer and editor for large datasets. Bioinformatics 30, 3276–3278.

Lau, C., Aubry, M., Musso, D., Teissier, A., Paulous, S., Desprès, P., de-Lamballerie, X., Pastorino, B., Cao-Lormeau, V.M., Weinstein, P., 2017. New evidence for endemic circulation of Ross River virus in the Pacific Islands and the potential for emergence. Int. J. Infect. Dis. 57, 73–76.

Lefort, V., Longueville, J.E., Gascuel, O., 2017. SMS: Smart Model Selection in PhyML. Mol. Biol. Evol. 34, 2422–2424.

Mackenzie, J., Lindsay, M.D.A., Coelen, R.J., Broom, A.K., Hall, R.A., Smith, D.W., 1994. Arboviruses causing human disease in the Australasian zoogeographic region. Arch. Virol. 136, 447–467.

Makau, D.N., Lycett, S., Michalska-smith, M., Paploski, I.A.D., Cheeran, M.C., Craft, M.E., Kao, R.R., Schroeder, D.C., Doeschl-wilson, A., Vanderwaal, K., 2022. Ecological and evolutionary dynamics of multi-strain RNA viruses. Nat. Ecol. Evol. 6, 1414–1422.

Marks, E.N., 1982. An atlas of common Queensland mosquitoes, revised. ed. Queensland Institute of Medical Research, Brisbane.

Marshall, I., Miles, J., 1984. Ross River virus and epidemic polyarthritis. In: KF, H. (Ed.), Current Topics in Vector Research. Praegcr Publications, New York, pp. 31–56.

Marshall, I., Woodroofe, G.M., Hirsch, S., 1982. Viruses recovered from mosquitoes and wildlife serum collected in the Murray Valley of South-eastern Australia, February 1974, during an epidemic of encephalitis. Aust. J. Exp. Biol. Med. Sci. 60, 457–470.

Martin, D.P., Murrell, B., Golden, M., Khoosal, A., Muhire, B., 2015. RDP4: Detection and analysis of recombination patterns in virus genomes. Virus Evol. 1, vev003.

Mcguinness, S.L., Lau, C.L., Leder, K., 2023. Co-circulation of Murray Valley encephalitis virus and Japanese encephalitis virus in south-eastern Australia. J. Travel Med. taad059.

Michie, A., Ernst, T., Chua, I.L.J., Lindsay, M.D.A., Neville, P.J., Nicholson, J., Jardine, A., Mackenzie, J.S., Smith, D.W., Imrie, A., 2020. Phylogenetic and Timescale Analysis of Barmah Forest Virus as Inferred from Genome Sequence Analysis. Viruses 12, 732.

Michie, A., Ernst, T., Pyke, A.T., Nicholson, J., Mackenzie, J.S., Smith, D.W., Imrie, A., 2023. Genomic Analysis of Sindbis Virus Reveals Uncharacterized Diversity within the Australasian Region, and Support for Revised SINV Taxonomy. Viruses 16, 7.

Michie, A., Mackenzie, J.S., Smith, D.W., Imrie, A., 2021. Genome sequence analysis of first Ross River virus isolate from Papua New Guinea indicates long-term, local evolution. Viruses 13, 482.

Moonen, J.P., Schinkel, M., van der Most, T., Miesen, P., van Rij, R.P., 2023. Composition and global distribution of the mosquito virome - A comprehensive database of insect-specific viruses. One Heal. 16, 100490.

Nahata, K.D., Bielejec, F., Monetta, J., Dellicour, S., Rambaut, A., Suchard, M.A., Baele, G., Lemey, P., 2022. SPREAD 4: online visualisation of pathogen phylogeographic reconstructions. Virus Evol. 8, veac088.

Nimmo, J.R., 1928. An unusual epidemic. Med. J. Aust. 1, 549–550.

NSW Government, N.E.H., 2023. NSW Arbovirus Surveillance and Mosquito Monitoring 2022–2023.

Nunes, M.R.T., Faria, N.R., Vasconcelos, J.M. De, Golding, N., Kraemer, M.U.G., Oliveira, L.F. De, Azevedo, R. do S. da S., Silva, D.E.A. da, Silva, E.V.P. da, Silva, S.P. da, Carvalho, V.L., Coelho, G.E., Cruz, A.C.R., Rodrigues, S.G., Vianez-Jr, J.L. da S.G., Nunes, B.T.D., Cardoso, J.F., Tesh, R.B., Hay, S.I., Pybus, O.G., Vasconcelos, P.F. da C., 2015. Emergence and potential for spread of Chikungunya virus in Brazil Emergence and potential for spread of Chikungunya virus in Brazil. BCM Med. 13, 102.

Ong, O.T.W., Skinner, E.B., Johnson, B.J., Old, J.M., 2021. Mosquito-Borne Viruses and Non-Human Vertebrates in Australia: A Review. Viruses 13, 265.

Patz, J.A., Norris, D.E., 2004. Land use change and human health. In: Geophysical Monograph Series. Blackwell Publishing Ltd, pp. 159–167.

Phillips, D., Sheridan, J., Aaskov, J., Murray, J., Wiemers, M., 1993. Epidemiology of Arbovirus Infection in Queensland, 1989-1992. Arbovirus Res. Aust. 6, 245–248.

Pickett, B.E., Sadat, E.L., Zhang, Y., Noronha, J.M., Squires, R.B., Hunt, V., Liu, M., Kumar, S., Zaremba, S., Gu, Z., Zhou, L., Larson, C.N., Dietrich, J., Klem, E.B., Scheuermann, R.H., 2012. ViPR: an open bioinformatics database and analysis resource for virology research. Nucleic Acids Res. 40, D593–D598.

Pollett, S., Fauver, J.R., Maljkovic Berry, I., Melendrez, M., Morrison, A., Gillis, L.D., Johansson, M.A., Jarman, R.G., Grubaugh, N.D., 2020. Genomic Epidemiology as a Public Health Tool to Combat Mosquito-Borne Virus Outbreaks. J. Infect. Dis. 221, S308– S318.

Powers, A., Huang, H., Roehrig, J., Strauss, E., Weaver, S., 2009. ICTV Ninth Report; Family Togaviridae.

Pronyk, P.M., Alwis, R. De, Rockett, R., Basile, K., Boucher, Y.F., Pang, V., Sessions, O., Getchell, M., Golubchik, T., Lam, C., Lin, R., Mak, T., Marais, B., Ong, R.T., Clapham, H.E., Wang, L., Cahyorini, Y., Polotan, F.G.M., Rukminiati, Y., Sim, E., Suster, C., Smith, G.J.D., Sintchenko, V., 2023. Advancing pathogen genomics in resource-limited settings. Cell Genomics 3, 100443.

Pyke, A.T., Shivas, M.A., Darbro, J.M., Onn, M.B., Johnson, P.H., Crunkhorn, A., Montgomery, I., Burtonclay, P., Jansen, C.C., Van Den Hurk, A.F., 2021. Uncovering the genetic diversity within the Aedes notoscriptus virome and isolation of new viruses from this highly urbanised and invasive mosquito. Virus Evol. 7, veab082.

Queensland Government, 2023. Land zones within Southeast Queensland bioregion [WWW Document]. URL https://apps.des.qld.gov.au/regional-ecosystems/landzones/?bioregion=12 (accessed 4.4.23).

Rambaut, A., 2014. FigTree: tree figure drawing tool version 1.4.3. University of Edinburgh, Edinburgh.

Rambaut, A., Drummond, A.J., Xie, D., Baele, G., Suchard, M.A., 2018. Posterior summarization in Bayesian phylogenetics using Tracer 1.7. Syst. Biol. 67, 901–904.

Rambaut, A., Lam, T.T., Carvalho, L.M., Pybus, O.G., 2016. Exploring the temporal structure of heterochronous sequences using TempEst (formerly Path-O-Gen). Virus Evol. 2, vew007.

Ritchie, S.A., Devine, G.J., Vazquez-Prokopec, G.M., Lenhart, A.E., Manrique-Saide, P., Scott, T.W., 2021. Insecticide-based approaches for dengue vector control. In: Koenraadt, C.J.M., Spitzen, J., Takken, W. (Eds.), Innovative Strategies for Vector Control – Progress in the Global Vector Control Response. Wageningen Academic Publishers, pp. 59–89.

Ritchie, S.A., Van Den Hurk, A.F., Zborowski, P., Kerlin, T.J., Banks, D., Walker, J.A., Lee, J.M., Montgomery, B.L., Smith, G.A., Pyke, A.T., Smith, I.L., 2007. Operational trials of remote mosquito trap systems for Japanese encephalitis virus surveillance in the Torres Strait, Australia. Vector-Borne Zoonotic Dis. 7, 497–506.

Robinson, J.T., Thorvaldsdóttir, H., Winckler, W., Guttman, M., Lander, E.S., Getz, G., Mesirov, J.P., 2011. Integrative genomics viewer. Nat. Biotechnol. 29, 24–26.

Russell, R.C., 1993. Mosquitoes and mosquito-borne disease in Southeastern Australia: A Guide to the Biology, Relation to Disease, Surveillance, Control and the Identification of Mosquitoes in Southeastern Australia, revised. ed. Westmead Hospital, Department of Medical Entomology, Sydney.

Russell, R.C., 2002. Ross River Virus: Ecology and Distribution. Annu. Rev. Entomol. 47, 1– 31.

Russell, R.C., Debenham, M.L., 1996. A colour photo atlas of mosquitoes of southeastern Australia. R.C. Russell, Sydney.

Selck, F.W., Adalja, A.A., Boddie, C.R., 2014. An estimate of the global health care and lost productivity costs of dengue. Vector-Borne Zoonotic Dis. 14, 824–826.

Shanks, G.D., 2019. Could Ross River Virus be the next Zika? J. Travel Med. 26, taz003.

Sikazwe, C., Neave, M.J., Michie, A., Mileto, P., Wang, J., Cooper, N., Levy, A., Imrie, A., Baird, R.W., Currie, B.J., Speers, D., Mackenzie, J.S., Smith, D.W., Williams, D.T., 2022. Molecular detection and characterisation of the first Japanese encephalitis virus belonging to genotype IV acquired in Australia. PLoS Negl. Trop. Dis. 16, e0010754.

Stanaway, J.D., Shepard, D.S., Undurraga, E.A., Halasa, Y.A., Coffeng, L.E., Brady, O.J., Hay, S.I., Bedi, N., Bensenor, I.M., Castañeda-Orjuela, C.A., Chuang, T.W., Gibney, K.B., Memish, Z.A., Rafay, A., Ukwaja, K.N., Yonemoto, N., Murray, C.J.L., 2016. The global burden of dengue: an analysis from the Global Burden of Disease Study 2013. Lancet Infect. Dis. 16, 712–723.

Stephenson, E.B., Murphy, A.K., Jansen, C.C., Peel, A.J., McCallum, H., 2019. Interpreting mosquito feeding patterns in Australia through an ecological lens: An analysis of blood meal studies. Parasites and Vectors 12, 156.

Stica, C.J., Barrero, R.A., Murray, R.Z., Devine, G.J., Phillips, M.J., Frentiu, F.D., 2022. Global Evolutionary History and Dynamics of Dengue Viruses Inferred from Whole Genome Sequences. Viruses 14, 703.

Streicker, D.G., Streicker, D.G., Turmelle, A.S., Vonhof, M.J., Kuzmin, I. V, Mccracken, G.F., Rupprecht, C.E., 2010. Establishment of Rabies Virus in Bats. Science (80-.). 329, 676– 679.

Suchard, M.A., Lemey, P., Baele, G., Ayres, D.L., Drummond, A.J., Rambaut, A., 2018. Bayesian phylogenetic and phylodynamic data integration using BEAST 1.10. Virus Evol. 4, vey016.

Toi, C.S., Webb, C.E., Haniotis, J., Clancy, J., Doggett, S.L., 2017. Seasonal activity, vector relationships and genetic analysis of mosquito-borne Stratford virus. PLoS One 12, e0173105.

van den Hurk, A.F., Hall-Mendelin, S., Johansen, C.A., Warrilow, D., Ritchie, S.A., 2012. Evolution of mosquito-based arbovirus surveillance systems in Australia. J. Biomed. Biotechnol. 2012, 325659.

van den Hurk, A.F., Hall-Mendelin, S., Townsend, M., Kurucz, N., Edwards, J., Ehlers, G., Rodwell, C., Moore, F.A., McMahon, J.L., Northill, J.A., Simmons, R.J., Cortis, G., Melville, L., Whelan, P.I., Ritchie, S.A., 2014. Applications of a sugar-based surveillance system to track arboviruses in wild mosquito populations. Vector-Borne Zoonotic Dis. 14, 66–73.

van Essen, P.H., Kemme, J.A., Ritchie, S.A., Kay, B.H., 1994. Differential responses of Aedes and Culex mosquitoes to octenol or light in combination with carbon dioxide in Queensland, Australia. Med. Vet. Entomol. 8, 63–67.

Victoria State Government, D. of H., 2023. Mosquito and mosquito-borne disease weekly report 2022/2023

Vieira, C.J. da S.P., Thies, S.F., da Silva, D.J.F., Kubiszeski, J.R., Barreto, E.S., Monteiro, H.A. de O., Mondini, A., São Bernardo, C.S., Bronzoni, R.V. de M., 2020. Ecological aspects of potential arbovirus vectors (Diptera: Culicidae) in an urban landscape of Southern Amazon, Brazil. Acta Trop. 202, 105276.

Vieira, C.J.S.P., Gyawali, N., Onn, M.B., Shivas, M.A., Shearman, D., Darbro, J.M., Wallau, G.L., van den Hurk, A.F., Frentiu, F.D., Skinner, E.B., Devine, G.J., 2024. Mosquito bloodmeals can be used to determine vertebrate diversity, host preference, and pathogen exposure in humans and wildlife. Res. Sq. [Preprint].

Vina-Rodriguez, A., Sachse, K., Ziegler, U., Chaintoutis, S.C., Keller, M., Groschup, M.H., Eiden, M., 2017. A Novel Pan-Flavivirus Detection and Identification Assay Based on RT-qPCR and Microarray. Biomed Res. Int. 2017, 4248756.

Volz, E.M., Koelle, K., Bedford, T., 2013. Viral Phylodynamics. Plos Comput. Biol. 9, e1002947.

Wash, R., Soria, C.D., 2015. True Blood: Dengue virus evolution. Nat. Rev. Microbiol. 13, 662.

WHO, W.H.O., 2023. Dengue and severe dengue.

Xu, G., Gao, T., Wang, Z., Zhang, J., Cui, B., Shen, X., Zhou, A., Zhang, Y., Zhao, J., Liu, H., Liang, G., 2023. Re-Emerged Genotype IV of Japanese Encephalitis Virus Is the Youngest Virus in Evolution. Viruses 15, 626.

Yakob, L., Hu, W., Frentiu, F.D., Gyawali, N., Hugo, L.E., Johnson, B., Lau, C., Furuya-Kanamori, L., Magalhaes, R.S., Devine, G., 2023. Japanese Encephalitis Emergence in Australia: The Potential Population at Risk. Clin. Infect. Dis. 76, 335–337.

Young, K.I., Medwid, J.T., Azar, S.R., Huff, R.M., Drumm, H., Coffey, L.L., Pitts, R.J., Buenemann, M., Vasilakis, N., Perera, D., Hanley, K.A., 2020. Identification of Mosquito Bloodmeals Collected in Diverse Habitats in Malaysian Borneo Using COI Barcoding. Trop. Med. Infect. Dis. 5, 51.

Zeghbib, S., Kemenesi, G., Jakab, F., 2023. The importance of equally accessible genomic surveillance in the age of pandemics. Biol. Futur. 74, 81–89.

Zhang, W., Yin, Q., Wang, H., Liang, G., 2023. The reemerging and outbreak of genotypes 4 and 5 of Japanese encephalitis virus. Front. Cell. Infect. Microbiol. 13, 1292693.

