## Supplementary data for "Long-term co-circulation of multiple arboviruses in southeast Australia revealed by xeno-monitoring and metatranscriptomics"

**Table S1:** The number of 2.3 mm zirconium silica beads (Daintree Scientific) and volume of DNA/RNA Shield (Zymo Research) that was added to each sample based on the number of mosquitoes in the pool.

| Mosquito no. | Bead no. | DNA/RNA Shield (uL) |
|---|---|---|
| **1 – 10** | 4 | 150 |
| **10 – 25** | 5 | 250 |
| **25 – 50** | 6 | 500 |
| **50 – 100** | 7 | 750 |
| **100 – 200** | 8 | 1200 |

**Table S2**: Genomes used for phylogenetic analysis (excel file).

**Table S3**: Tempest results to examine the temporal signal of data sets prior to phylogenetic molecular clock analyses.

| Tempest analysis | Quantity of sequences | Period | Correlation coefficient | $R^2$ |
|---|---|---|---|---|
| ARGV G3C clade currently circulating in Australia | 11 | 2004–2022 | 0.8817 | 0.7774 |
| All ARGV complete sequences isolated from Australasia | 66 | 1960–2022 | 0.8623 | 0.7436 |
| All ARGV complete sequences isolated from Australasia + China | 67 | 1960–2022 | 0.5689 | 0.3237 |
| BFV G3B subclade currently circulating in Australia | 11 | 2013–2022 | 0.9194 | 0.8453 |
| BFV complete sequences isolated from Australia | 38 | 1974–2022 | 0.8634 | 0.7455 |
| BFV complete sequences isolated from Australia and PNG | 39 | 1974–2022 | 0.6546 | 0.4285 |
| RRV G4B subclade currently circulating in Australia | 30 | 2008–2022 | 0.9758 | 0.9521 |
| RRV complete sequences isolated from Australia, PICTs and PNG | 141 | 1959–2022 | 0.8057 | 0.6492 |
| STRV most recent genomes isolated in Australia | 9 | 2006–2022 | 0.5795 | 0.3358 |
| All STRV whole genomes isolated in Australia | 11 | 1961–2022 | 0.949 | 0.9005 |

All data sets exhibit a positive correlation between genetic divergence and sampling time and should be suitable for phylogenetic molecular clock analysis in BEAST or other programs. However, datasets in red were not used.

**Table S4**: Path sampling (PS) and Stepping-stone (SS) log marginal likelihood analysis of different molecular clock and demographic models on different alignments.

| All 66 ARGV whole-genome sequences isolated in Australia from 1960–2002 | | | |
|---|---|---|---|
| **Clock** | **Demographic prior** | **PS** | **SS** |
| UCLN | Skyline | -29116.212960828874 | -29116.085969376585 |
| UCLN | Constant | -29112.88377327311 | -29112.578372824344 |
| Strict | Skyline | -29188.085694987047 | -29188.581916119 |
| Strict | Constant | -29186.929182892934 | -29186.762066982345 |
| All 39 BFV whole-genome sequences isolated in Australia and PNG from 1974–2002 | | | |
| **Clock** | **Clock** | **Clock** | **Clock** |
| UCLN | Skyline | -20766.8856632555 | -20767.441244682843 |
| UCLN | Constant | -20763.844254276974 | -20763.939892189726 |
| Strict | Skyline | -20760.469291502788 | -20760.738569310015 |
| Strict | Constant | -20760.984178069684 | -20760.889966051578 |
| All 141 RRV whole-genome sequences isolated in Australasia from 1959–2022 | | | |
| **Clock** | **Demographic prior** | **PS** | **SS** |
| UCLN | Skyline | -33562.03697370134 | -33563.42913163195 |
| UCLN | Constant | -33560.64553727518 | -33561.436506728794 |
| Strict | Skyline | -33674.307081194456 | -33674.998528977194 |
| Strict | Constant | -33676.09783559379 | -33676.79366004619 |
| All 11 STRV whole-genome sequences isolated in Australia from 1961–2022 | | | |
| **Clock** | **Demographic prior** | **PS** | **SS** |
| UCLN | Skyline | -17870.398499805175 | -17870.482838006646 |
| UCLN | Constant | -17868.346193924048 | -17868.3209418113 |
| Strict | Skyline | -17862.53541325153 | -17862.593564144074 |
| Strict | Constant | -17860.29563175868 | -17860.226743438852 |

The better models are highlighted in yellow.

**Table S5:** Mosquito species collected across Brisbane sites, from March 2021 to May 2022.

| Species | Banyo (%) | D% | Bracken Ridge (%) | D% | Corinda (%) | D% | Hemmant (%) | D% | Indooroopilly (%) | D% | Abundance (%) | D% |
|---|---|---|---|---|---|---|---|---|---|---|---|---|
| *Aedes aculeatus* | 9 (<0.1) | Rr | 3 (<0.1) | Rr | 1 (<0.1) | Rr | 0 | | 1 (0.1) | Rr | 14 (<0.1) | Rr |
| *Ae. alboannulatus* | 0 | | 0 | | 0 | | 1 (4.5) | Rr | 0 | | 1 (<0.1) | Rr |
| *Ae. alboscutellatus* | 4 (<0.1) | Rr | 0 | | 33 (0.4) | Rr | 0 | | 0 | | 37 (0.1) | Rr |
| *Ae. alternans* | 38 (0.1) | Rr | 6 (<0.1) | Rr | 0 | | 0 | | 0 | | 44 (0.1) | Rr |
| *Ae. kochi* | 0 | | 0 | | 0 | | 0 | | 2 (0.2) | Rr | 2 (<0.1) | Rr |
| *Ae. lineatopennis* | 11 (<0.1) | Rr | 4 (<0.1) | Rr | 4 (0.1) | Rr | 0 | | 0 | | 19 (<0.1) | Rr |
| *Ae. notoscriptus** | 71 (0.3) | Rr | 37 (0.2) | Rr | 87 (1.1) | Ev | 0 | | 33 (3.5) | Sd | 228 (0.4) | Rr |
| ***Ae. procax**** | **3,005 (11.7)** | E | 382 (1.9) | Ev | 862 (11.2) | E | 2 (9.1) | D | **128 (13.6)** | E | **4,379 (8.1)** | D |
| *Ae. theobaldi* | 0 | | 0 | | 0 | | 0 | | 1 (0.1) | Rr | 1 (<0.1) | Rr |
| ***Ae. vigilax**** | **2,170 (8.4)** | D | **1,346 (6.8)** | D | 17 (0.2) | Rr | **11 (50.0)** | E | 22 (2.3) | Sd | 3,566 (6.6) | D |
| *Ae. vittiger* | 37 (0.1) | Rr | 9 (<0.1) | Rr | 1 (<0.1) | Rr | 0 | | 74 (7.9) | D | 121 (0.2) | Rr |
| *Anopheles annulipes** | 25 (0.1) | Rr | 25 (0.1) | Rr | 4 (0.1) | Rr | 0 | | 5 (0.5) | Rr | 59 (0.1) | Rr |
| *An. atratipes* | 2 (<0.1) | Rr | 14 (0.1) | Rr | | | 0 | | 1 (0.1) | Rr | 17 (<0.1) | Rr |
| *An. bancroftii* | 15 (0.1) | Rr | 1 (<0.1) | Rr | 148 (1.9) | Ev | 0 | | 1 (0.1) | Rr | 165 (0.3) | Rr |
| *Coquillettidia linealis** | 206 (0.8) | Rr | 1,293 (6.5) | D | 7 (0.1) | Rr | 0 | | 2 (0.2) | Rr | 1,508 (2.8) | Sd |
| *Cq. xanthogaster** | 629 (2.4) | Sd | 782 (4.0) | Sd | 420 (5.5) | D | 0 | | 0 | | 1,831 (3.4) | Sd |
| ***Culex annulirostris**** | **16,186 (62.8)** | E | **9,354 (47.3)** | E | **2,841 (37.1)** | E | **3 (13.6)** | E | **587 (62.4)** | E | **2,8971 (53.5)** | E |
| ***Cx. orbostiensis*** | 2,083 (8.1) | D | **4,206 (21.3)** | E | **1,805 (23.5)** | E | 1 (4.5) | Sd | 2 (0.2) | Rr | **8,097 (14.9)** | E |
| *Cx. quinquefasciatus** | 1 (<0.1) | Rr | 1 (<0.1) | Rr | 1 (<0.1) | Rr | 2 (9.1) | D | 0 | | 5 (<0.1) | Rr |
| *Cx. sitiens** | 495 (1.9) | Ev | 394 (2.0) | Ev | 231 (3.0) | Sd | 2 (9.1) | D | 3 (0.3) | Rr | 1,125 (2.1) | Sd |
| ***Mansonia uniformis**** | 52 (0.2) | Rr | 40 (0.2) | Rr | **963 (12.6)** | E | 0 | | 1 (0.1) | Rr | 1,056 (1.9) | Ev |
| *Mimomiya elegans* | 5 (<0.1) | Rr | 9 (<0.1) | Rr | 10 (0.1) | Rr | 0 | | 0 | | 24 (<0.1) | Rr |
| *Uranotenea nivipes* | 6 (<0.1) | Rr | | | 2 (<0.1) | Rr | 0 | | 0 | | 8 (<0.1) | Rr |
| *Ur. pygmaea* | 2 (<0.1) | Rr | 52 (0.3) | Rr | 1 (<0.1) | Rr | 0 | | 0 | | 55 (0.1) | Rr |
| ***Verralinea funerea**** | 668 (2.6) | Sd | 823 (4.2) | Sd | 25 (0.3) | Rr | 0 | | **77 (8.2)** | D | 1,593 (2.9) | Sd |
| *Ve.* Marks sp52 | 51 (0.2) | Rr | 1,008 (5.1) | D | 205 (2.7) | Sd | 0 | | 0 | | 1,264 (2.3) | Sd |
| Total abundance (%) | 25,771 (47.6) | | 19,789 (36.5) | | 7,668 (14.2) | | 22 (<0.1) | | 940 (1.7) | | 54,190 (100) | |
| Species richness | 23 | | 21 | | 21 | | 7 | | 16 | | 26 | |
| Number of collections | 8 | | 5 | | 5 | | 1 | | 2 | | 21 | |
| Number of traps | 23 | | 13 | | 17 | | 1 | | 4 | | 58 | |

* Species demonstrated as competent RRV vectors in laboratory studies (Russell, 2002).

Most abundant species in bold.

D%: Dominance – E: eudominant (D > 10%); D: dominant (D > 5–10%); Sd: subdominant (D > 2–5%); Ev: eventual (D > 1–2%); Rr: rare (D < 1%)

**Table S6:** Web addresses for visualisation of animation over time of maximum clade credibility (MCC) annotated trees summarising either a discrete or continuous phylogeographic reconstruction.

| **Phylogeographic analysis** | **Web address** |
|---|---|
| SINV-like: G3C clade currently circulating in Australia (11 sequences) continuous MCC tree | https://view.spreadviz.org/?output=3bfb6806-8773-43a9-aeb8-b544a973baaf/b928ef8d-8a42-4d78-8a2f-9d28af67f5bb.json&maps=AU |
| SINV-like: G3C clade currently circulating in Australia (11 sequences) discrete MCC tree | https://view.spreadviz.org/?output=3bfb6806-8773-43a9-aeb8-b544a973baaf/3d816026-5712-495f-82d8-35bbf4e828e5.json&maps=AU |
| BFV: G3B clade currently circulating in Australia (11 sequences) continuous MCC tree | https://view.spreadviz.org/?output=3bfb6806-8773-43a9-aeb8-b544a973baaf/fe533490-3bb6-4873-bfb5-a6ab3f41e3bd.json&maps=AU |
| BFV: G3B clade currently circulating in Australia (11 sequences) discrete MCC tree | https://view.spreadviz.org/?output=3bfb6806-8773-43a9-aeb8-b544a973baaf/37a5c2f0-9f7b-4ca4-a772-72ca3a062166.json&maps=AU |
| RRV: G4B clade currently circulating in Australia (23 sequences) continuous MCC tree | https://view.spreadviz.org/?output=3bfb6806-8773-43a9-aeb8-b544a973baaf/267fda95-2c8a-423e-94d8-4c69c05fed6b.json&maps=AU |
| RRV: G4B clade currently circulating in Australia (30 sequences) discrete MCC tree | https://view.spreadviz.org/?output=3bfb6806-8773-43a9-aeb8-b544a973baaf/0d24c743-1c25-46c2-b7dc-39b83fb83b2c.json&maps=AU |
| STRV: all whole genomes isolated in Australia (11 sequences) continuous MCC tree | https://view.spreadviz.org/?output=3bfb6806-8773-43a9-aeb8-b544a973baaf/b34bb117-24c1-4f34-aa4f-4f60efe8caf5.json&maps=AU |
| STRV: all whole genomes isolated in Australia (11 sequences) discrete MCC tree | https://view.spreadviz.org/?output=3bfb6806-8773-43a9-aeb8-b544a973baaf/18e84ce3-4ebb-4845-8a2e-03b4d9f12422.json&maps=AU |

**a) SINV-like G3C clade currently circulating in Australia continuous and discrete maps**

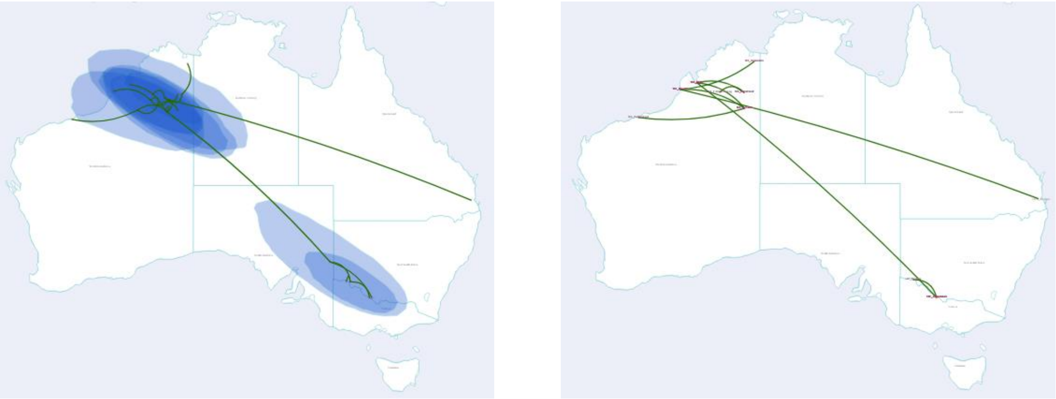

**b) BFV G3B clade currently circulating in Australia continuous and discrete maps**

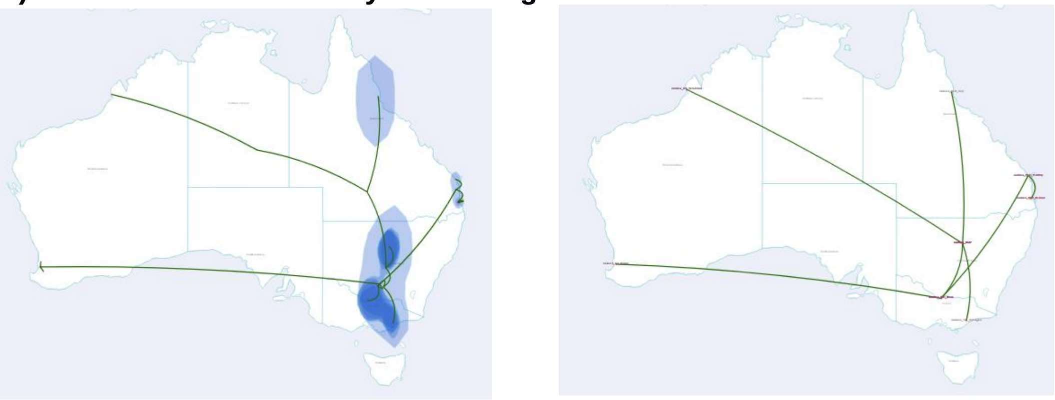

**c) RRV G4B clade currently circulating in Australia continuous and discrete maps**

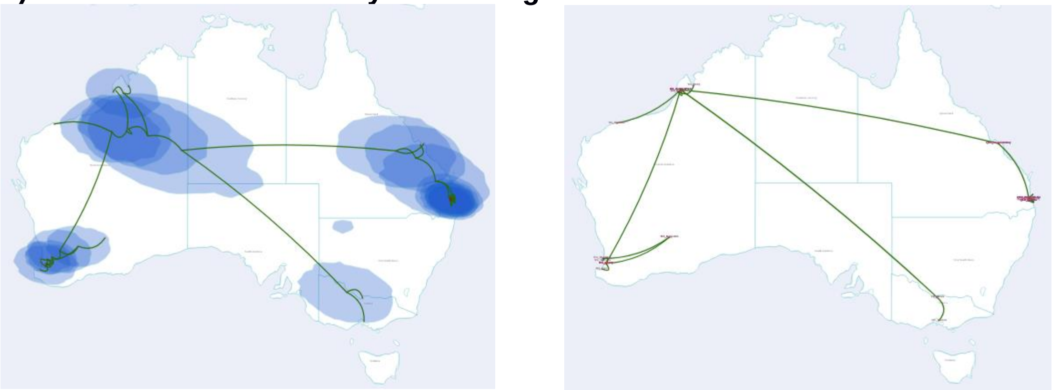

**d) All complete genomes of STRV continuous and discrete maps**

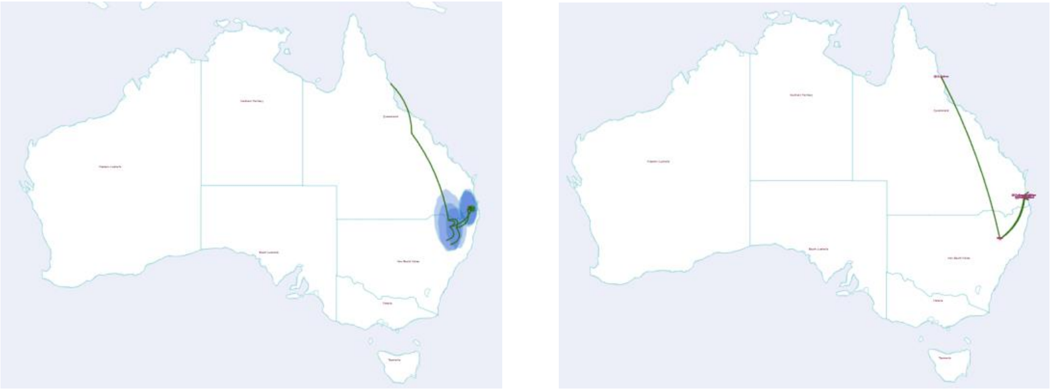

**Figure S1.** Maps produced using SPREAD 4 web server.

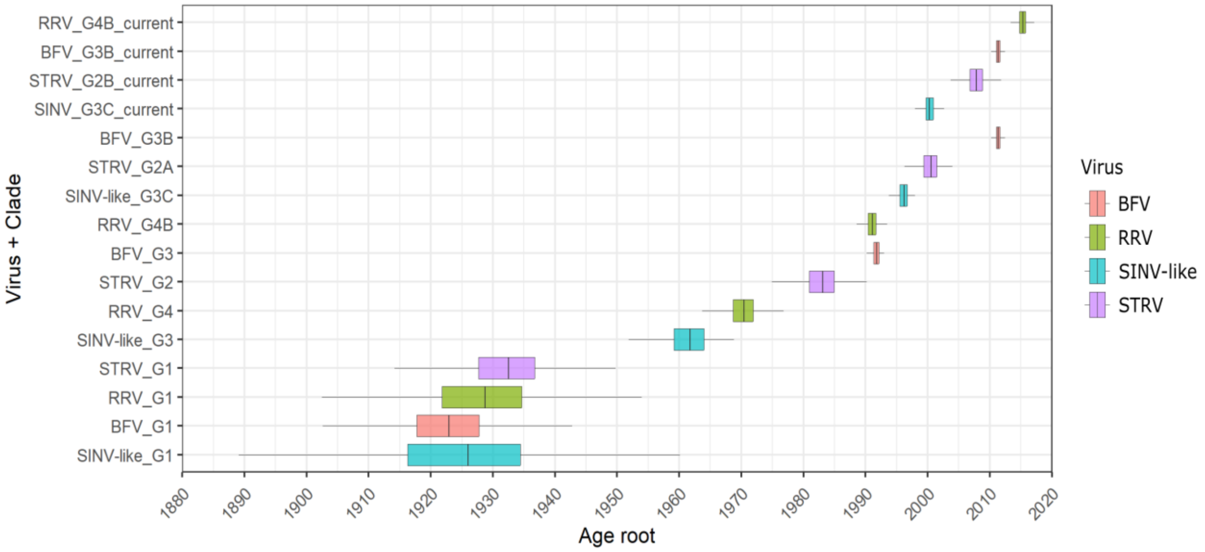


**Figure S2.** Temporal analysis of arboviruses clades.

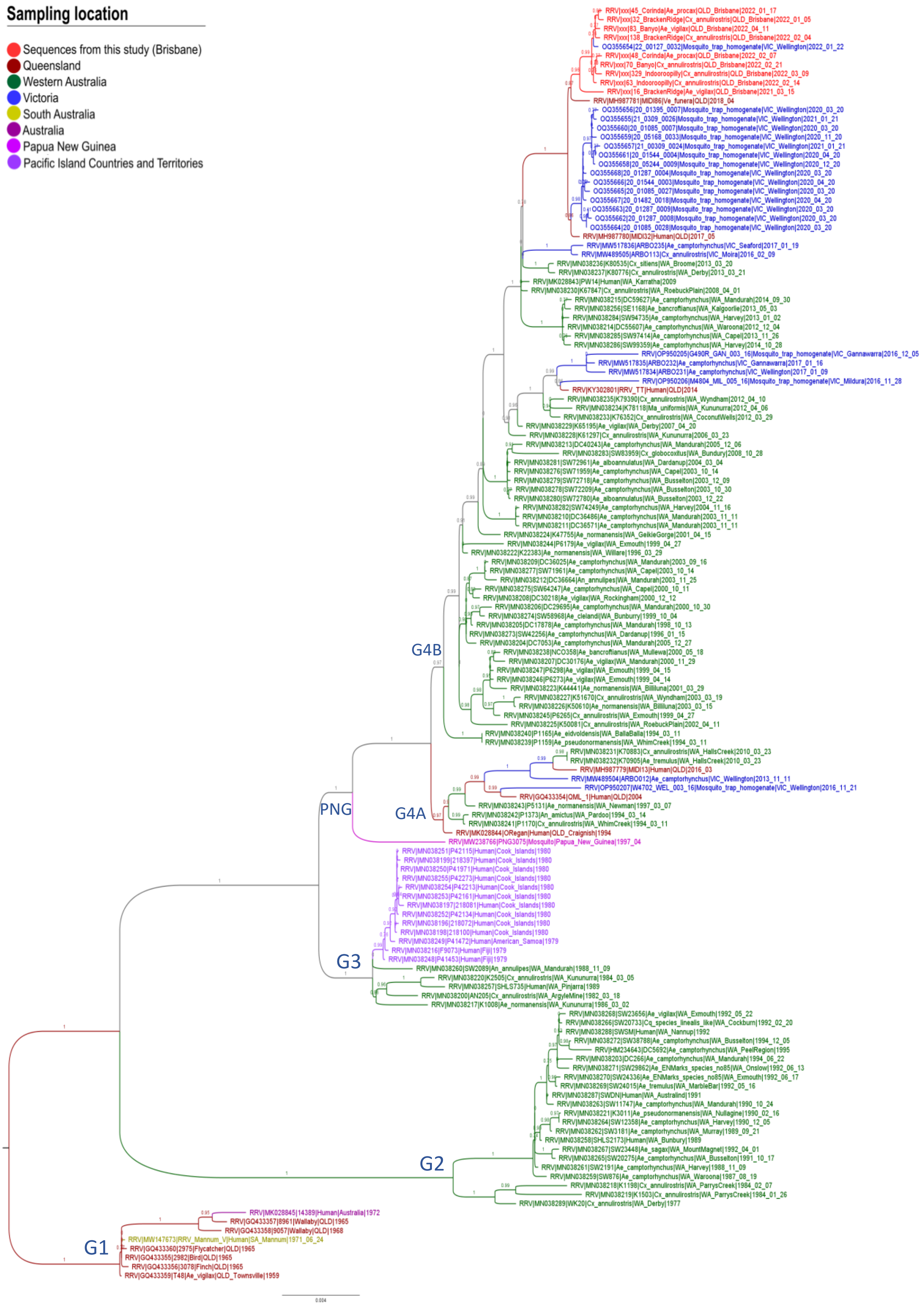


**Figure S3.** Maximum likelihood (ML) phylogeny reconstruction of RRV including all available whole-genome and draft genomes >10 kb (a total of 141 sequences). aLRT SH-like branch support of key clades are presented above nodes.

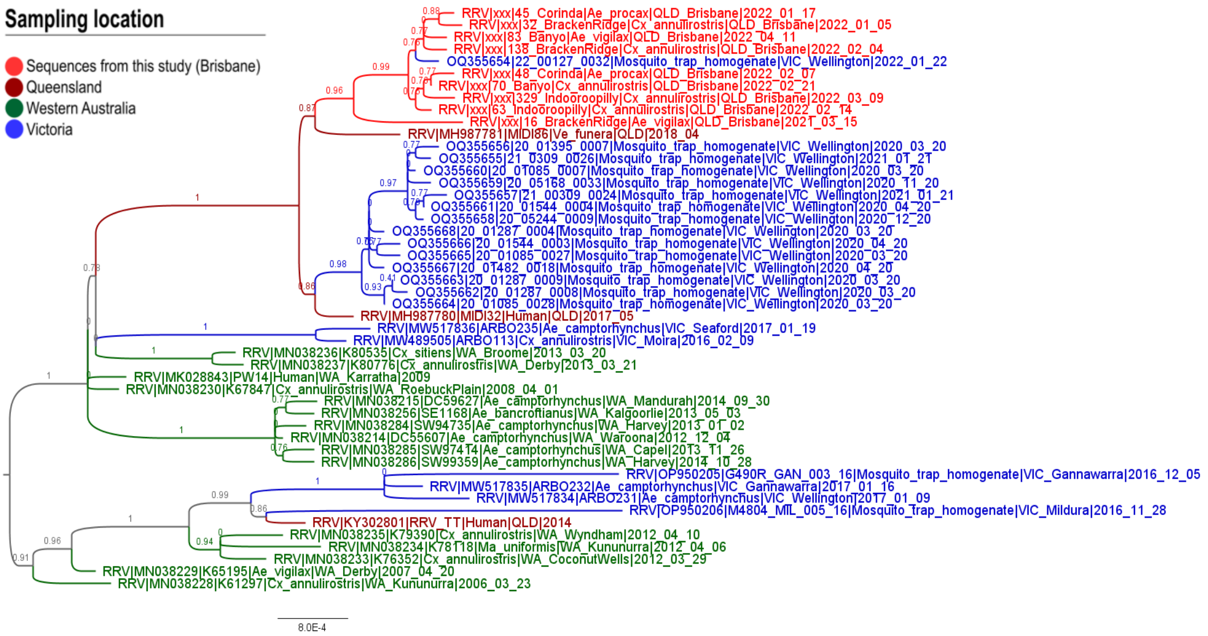


**Figure S4**. ML phylogeny reconstruction of RRV G4B genotype currently circulating in Australia including all available whole-genome and draft genomes >10 kb (a total of 48 sequences). aLRT SH-like branch support of key clades are presented above nodes.

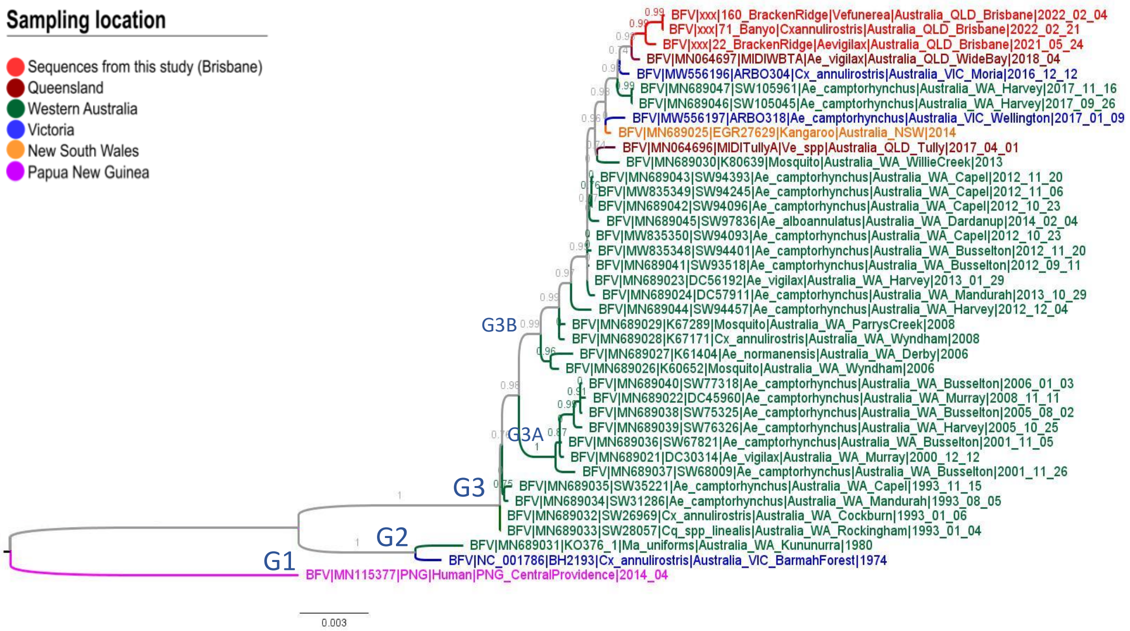


**Figure S5.** ML phylogeny reconstruction of BFV including all available whole-genome and draft genomes >10 kb (a total of 39 sequences). aLRT SH-like branch support of key clades are presented above nodes.

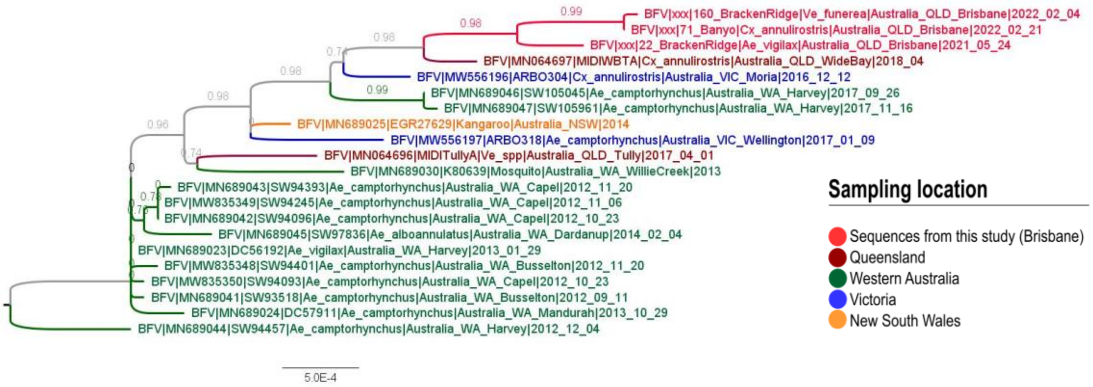


**Figure S6**. ML phylogeny reconstruction of BFV G3B genotype currently circulating in Australia including all available whole-genome and draft genomes >10 kb (a total of 21 sequences). aLRT SH-like branch support of key clades are presented above nodes.

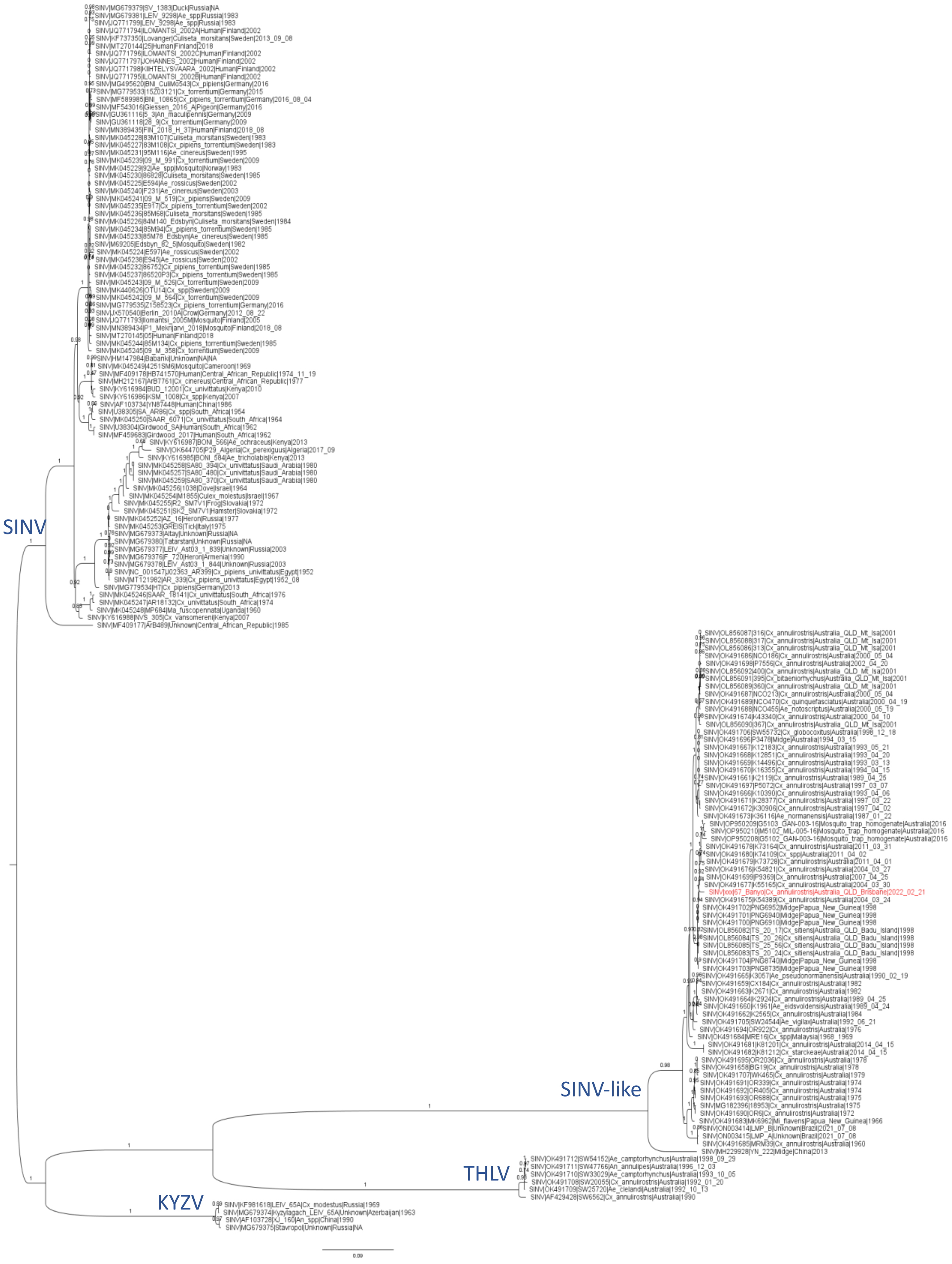


**Figure S7.** ML phylogeny reconstruction of SINV including all available whole-genome and draft genomes >10 kb (a total of 161 sequences). aLRT SH-like branch support of key clades are presented above nodes. Virus nomenclature includes the GenBank accession numbers, strain name, location of collection, species of origin, and the year of sampling. Red tip represents the new sequence obtained in this study. SINV – Sindbis virus; SINV-like – Argyle virus; KYZV – Kyzylagach virus; THLV – Thomson's Lake virus.

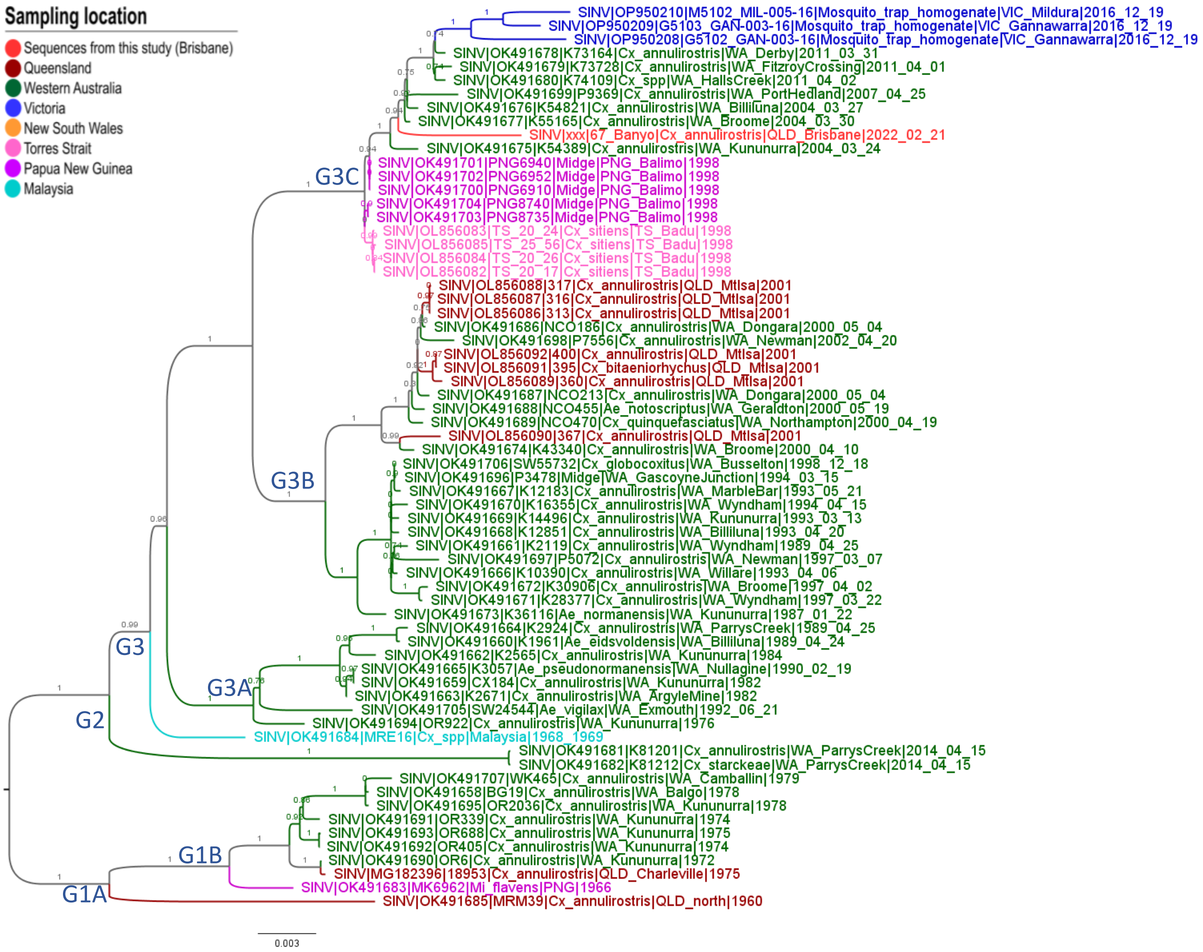


**Figure S8.** ML phylogeny reconstruction of SINV-like including all available whole-genome and draft genomes >10 kb (a total of 66 sequences). aLRT SH-like branch support of key clades are presented above nodes.

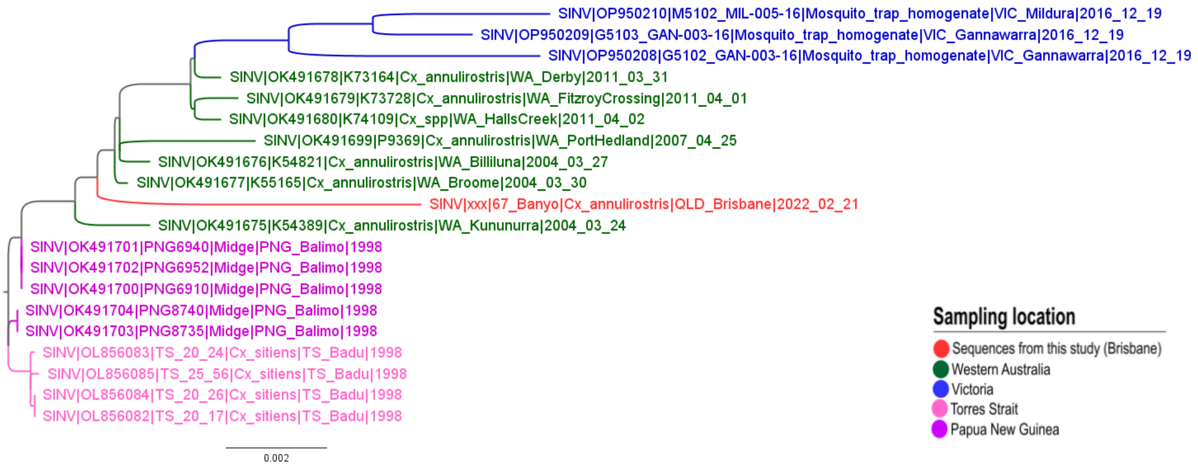


**Figure S9.** ML phylogeny reconstruction of SINV-like G3C genotype currently circulating in Australia including all available whole-genome and draft genomes >10 kb (a total of 20 sequences). aLRT SH-like branch support of key clades are presented above nodes.

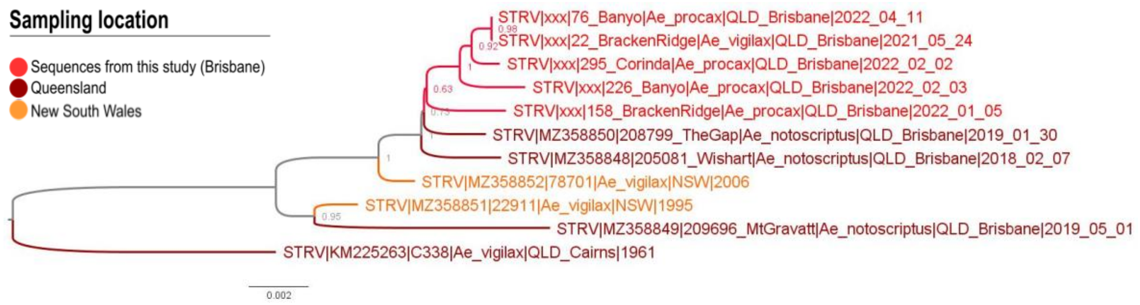


**Figure S10.** ML phylogeny reconstruction of STRV including all available whole-genome and draft genomes >9 kb (a total of 11 sequences). aLRT SH-like branch support of key clades are presented in front of the nodes.

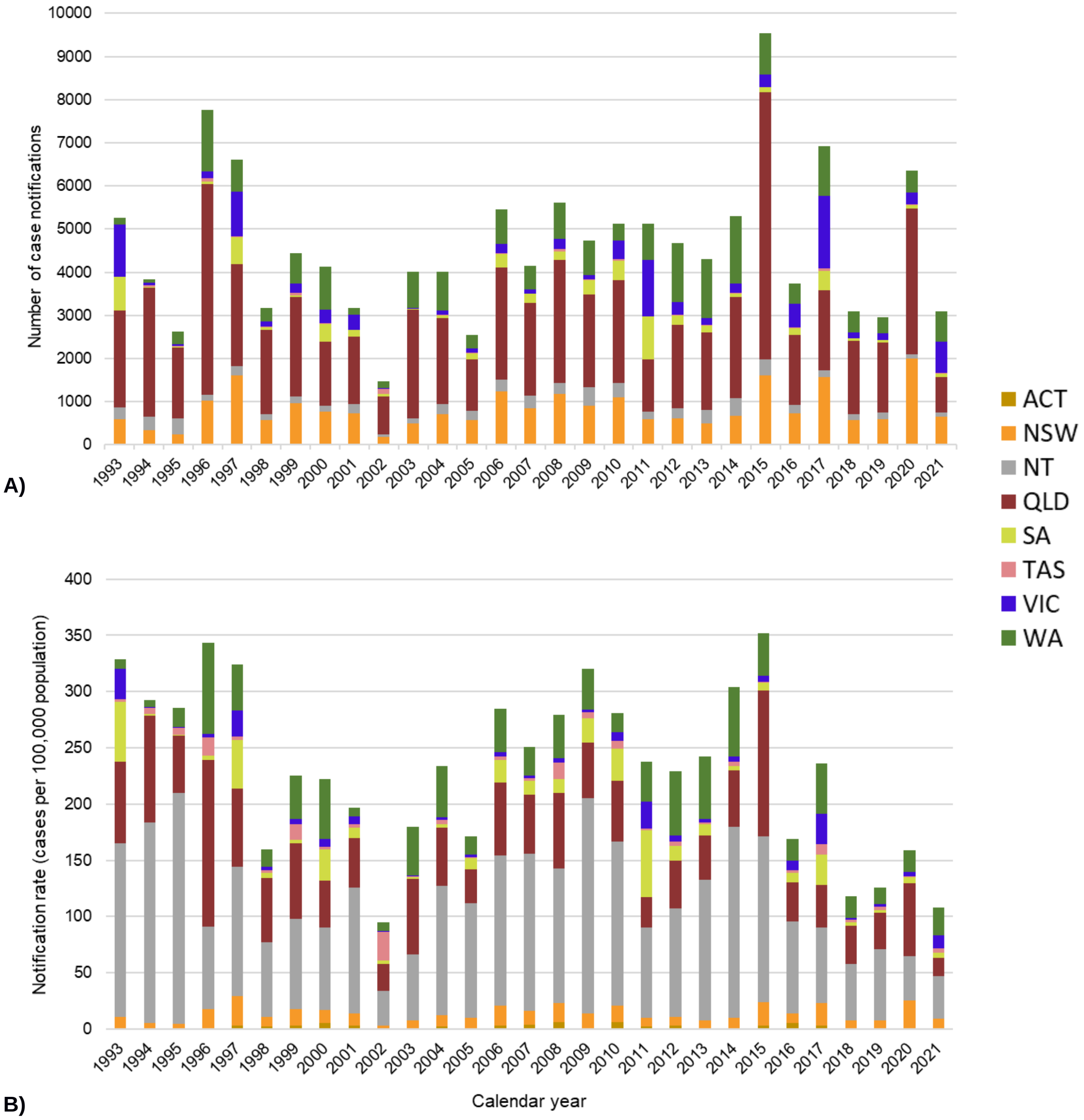


**Figure S11.** The epidemiology of Ross River virus (RRV) in Australia from 1993 to 2021. Graph A illustrates the number of notified RRV cases per calendar year within each state. Graph B illustrates the incidence rate of RRV infections per 100,000 population per calendar year within each state. The data used for these graphs was sourced from the National Notifiable Disease Surveillance System (NNDSS) in May 2023.

Australian states: ACT – Australian Capital Territory; NSW – New South Wales; NT – Northern Territory; QLD – Queensland; SA – South Australia; TAS – Tasmania; VIC – Victoria; WA – Western Australia.

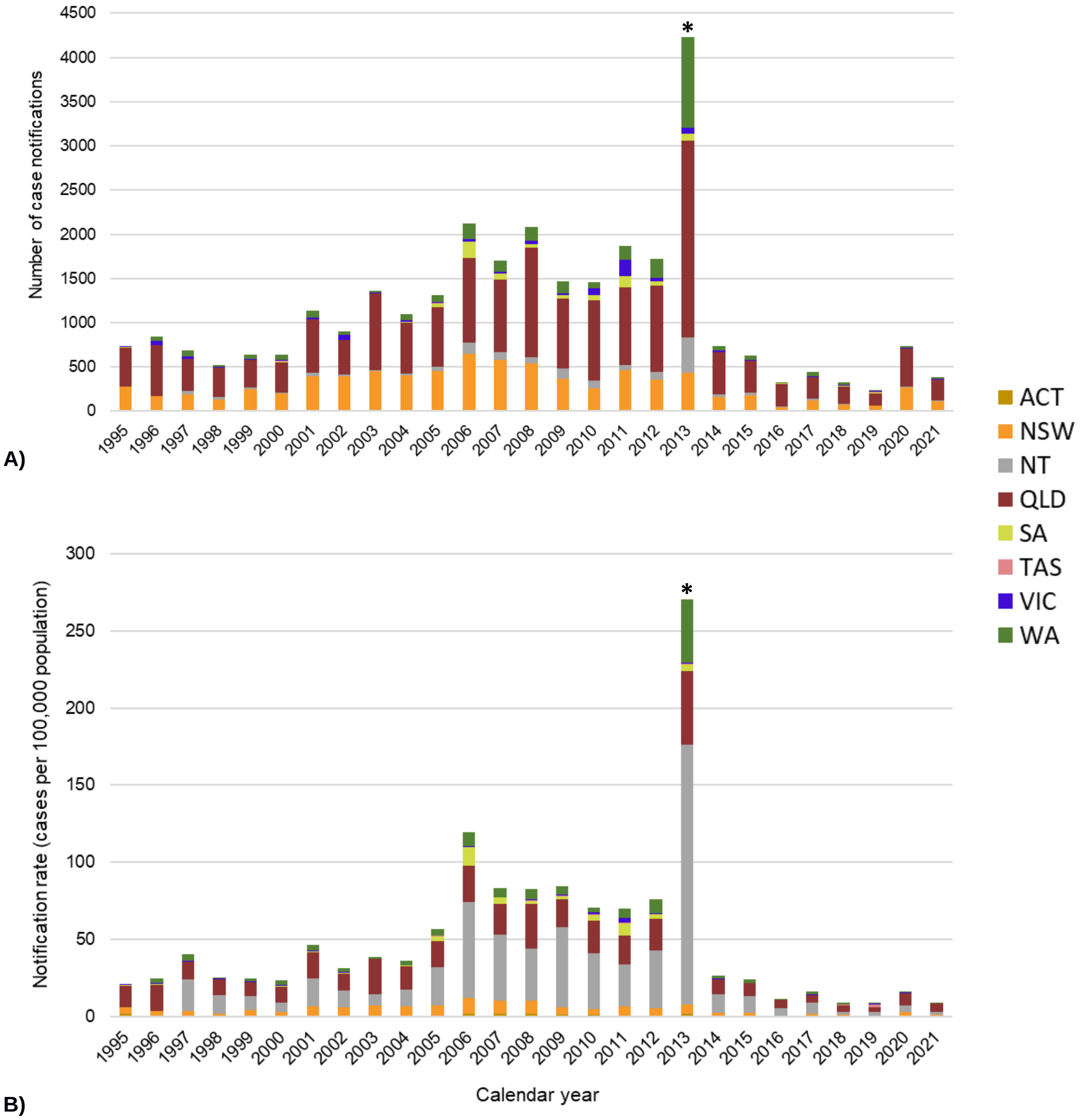


**Figure S12.** The epidemiology of Barmah Forest virus (BFV) in Australia from 1995 to 2021. Graph A illustrates the number of notified BFV cases per calendar year within each state. Graph B illustrates the incidence rate of BFV infections per 100,000 population per calendar year within each state. Notification from the year 2013 (*) should be discounted. This particular year experienced a high number of false-positive case reports due to the widespread use of the faulty Alere PanBio BFV IgM ELISA kit. The data used for these graphs was sourced from the National Notifiable Disease Surveillance System (NNDSS) in May 2023.

Australian states: ACT – Australian Capital Territory; NSW – New South Wales; NT – Northern Territory; QLD – Queensland; SA – South Australia; TAS – Tasmania; VIC – Victoria; WA – Western Australia.

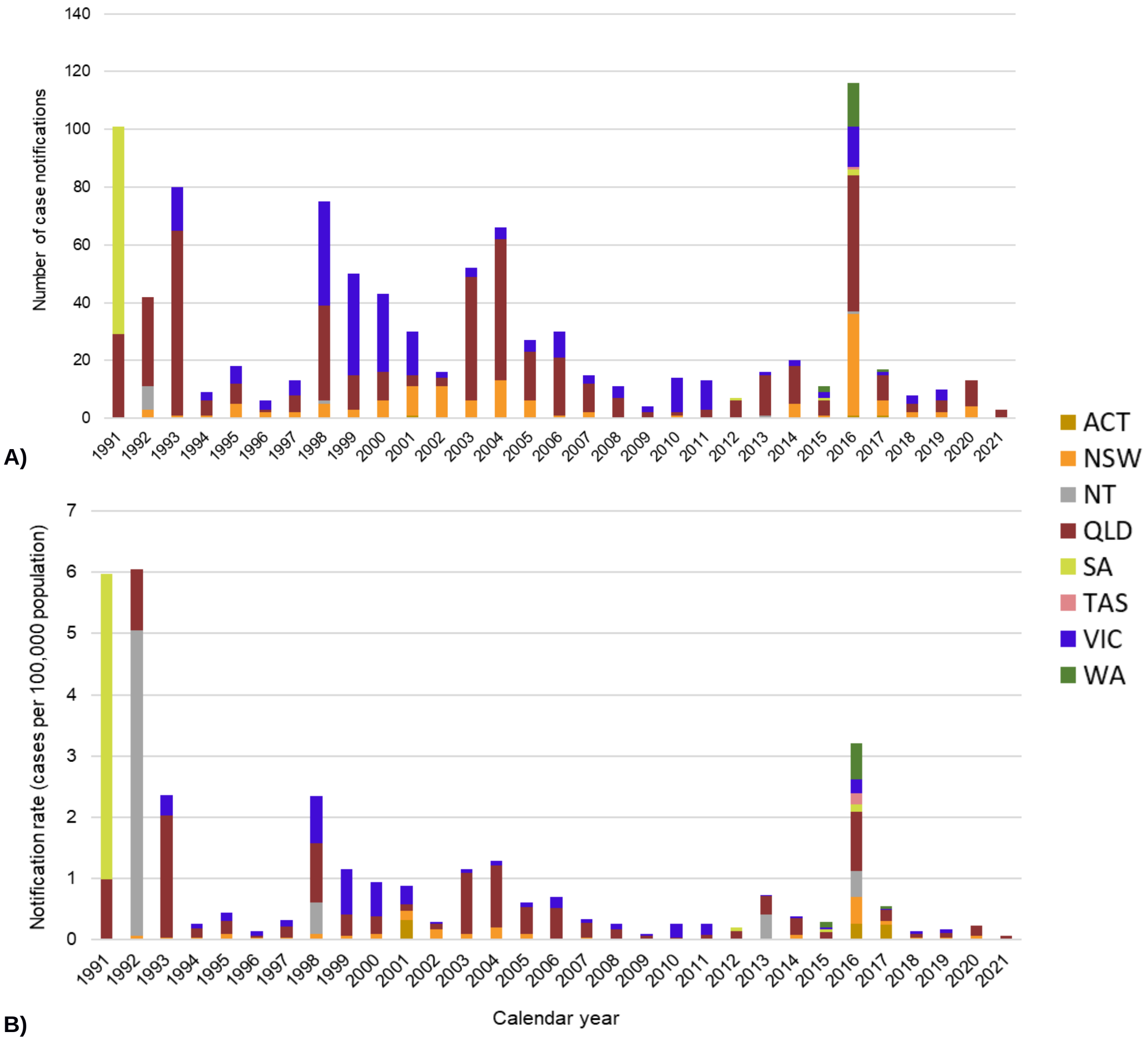


**Figure S13.** The epidemiology of unspecified Flavivirus infections in Australia from 1991 to 2021. Graph A illustrates the number of Flavivirus infections notified per calendar year within each state. Graph B illustrates the incidence rate of Flavivirus infections per 100,000 population per calendar year within each state. The data used for these graphs was sourced from the National Notifiable Disease Surveillance System (NNDSS) in May 2023.

Australian states: ACT – Australian Capital Territory; NSW – New South Wales; NT – Northern Territory; QLD – Queensland; SA – South Australia; TAS – Tasmania; VIC – Victoria; WA – Western Australia.

**Table S7:** Virus detections from whole trap grinds in the 2022-2023 surveillance season.

| Virus detected | Australian state | | | TOTAL |
|---|---|---|---|---|
| | NSW | VIC | SA | |
| **RRV** | 4 | 21 | 19 | **44** |
| **BFV** | 9 | 11 | 13 | **33** |
| **WNV/KUN** | 2 | 8 | 2 | **12** |
| **MVEV** | 18 | 48 | 11 | **77** |
| **STRV** | 7 | Not tested | Not tested | **7** |
| **EHV** | 6 | Not tested | Not tested | **6** |
| **TOTAL** | **46** | **88** | **45** | **179** |

Arboviruses: RRV – Ross river virus; BFV – Barmah forest virus; WNV/KUNV – West Nile virus / Kunjin strain; MVEV – Murray valley encephalitis; STRV – Stratford virus; EHV – Edge Hill virus.
Australian states: NSW – New South Wales; SA – South Australia; VIC – Victoria.

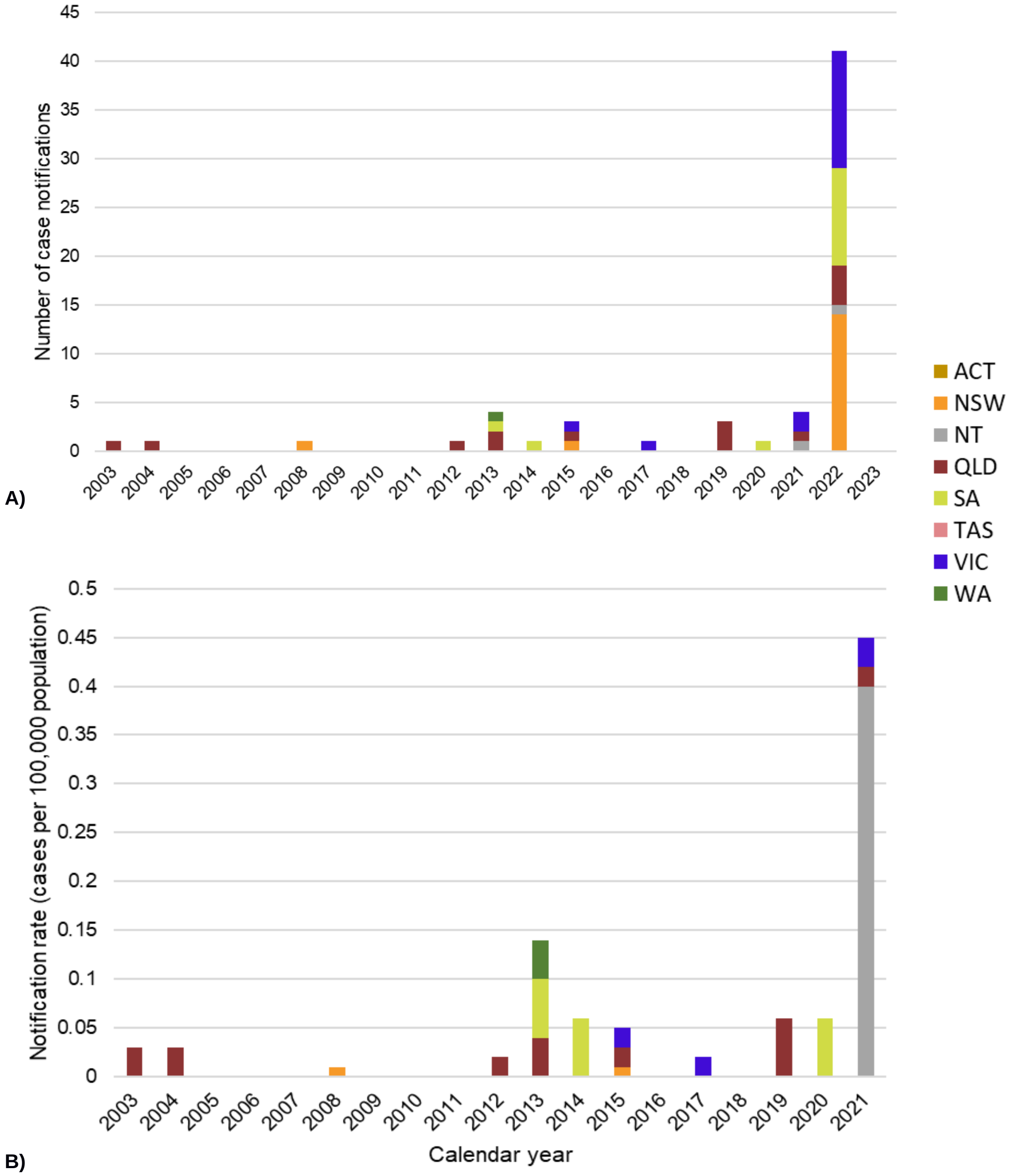


**Figure S14.** The epidemiology of Japanese Encephalitis virus (JEV) in Australia from 2003 to 2021/23. Graph A illustrates the number of JEV notified cases per calendar year within each state. Graph B illustrates the incidence rate of JEV infections per 100,000 population per calendar year within each state. The data used for these graphs was sourced from the National Notifiable Disease Surveillance System (NNDSS) in May 2023.

Australian states: ACT – Australian Capital Territory; NSW – New South Wales; NT – Northern Territory; QLD – Queensland; SA – South Australia; TAS – Tasmania; VIC – Victoria; WA – Western Australia.

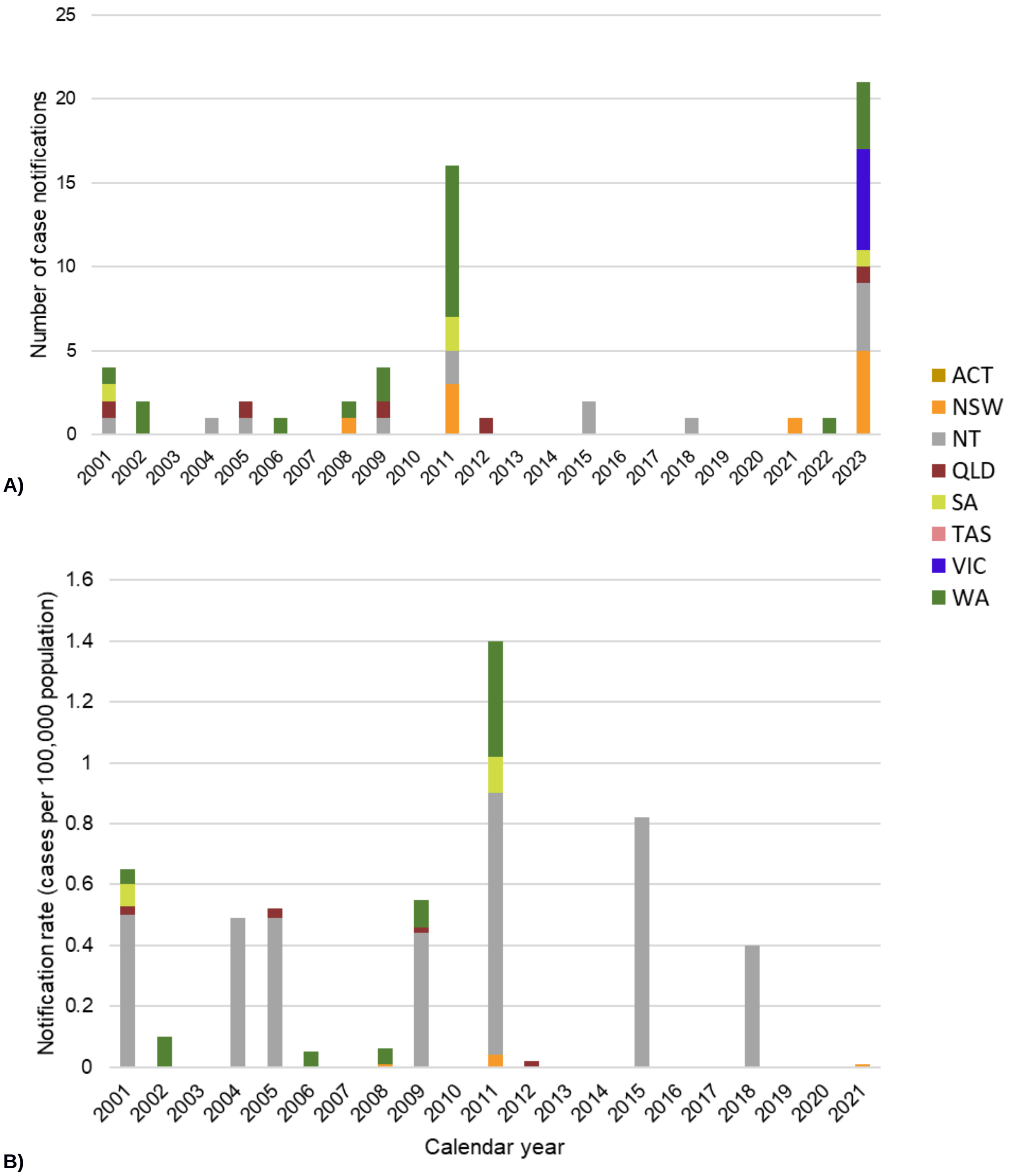


**Figure S15.** The epidemiology of Murray Valley Encephalitis virus (MVEV) in Australia from 2001 to 2021/23. Graph A illustrates the number of MVEV notified cases per calendar year within each state. Graph B illustrates the incidence rate of MVEV infections per 100,000 population per calendar year within each state. The data used for these graphs was sourced from the National Notifiable Disease Surveillance System (NNDSS) in May 2023.

Australian states: ACT – Australian Capital Territory; NSW – New South Wales; NT – Northern Territory; QLD – Queensland; SA – South Australia; TAS – Tasmania; VIC – Victoria; WA – Western Australia.

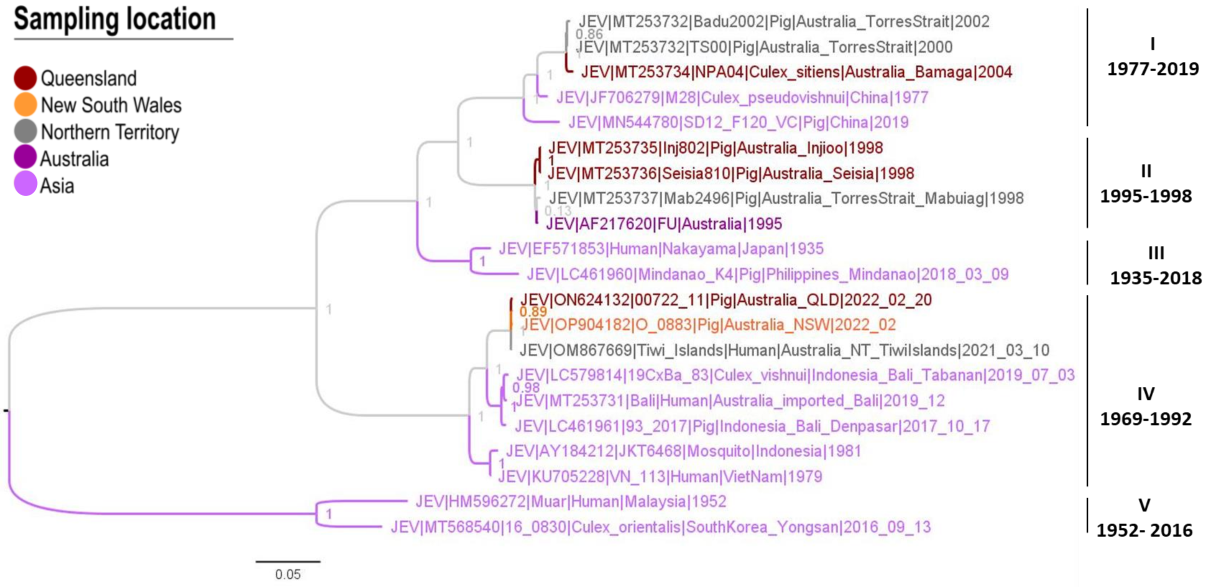


**Figure S16.** ML phylogeny reconstruction of JEV including all available Australian whole-genome and draft genomes >9 kb (a total of **ten** sequences from 1995 to 2022) plus some Asian representative genomes from all genotypes of JEV. aLRT SH-like branch support of key clades are presented in front of the nodes. Genotype in red is the one that recently emerged in Australia.

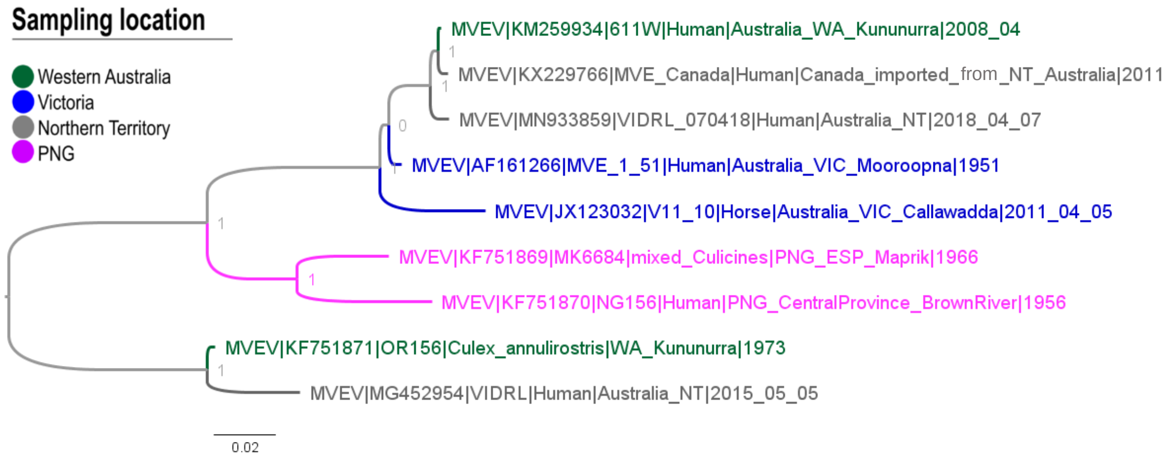


**Figure S17.** ML phylogeny reconstruction of MVEV including all available whole-genome and draft genomes >9 kb (a total of **nine** sequences, **seven** from Australia from 1951 to 2018). aLRT SH-like branch support of key clades are presented in front of the nodes.
